# Cellular segregation in co-cultures driven by differential adhesion and contractility on distinct time scales

**DOI:** 10.1101/2022.05.23.492966

**Authors:** Mark Skamrahl, Justus Schünemann, Markus Mukenhirn, Hongtao Pang, Jannis Gottwald, Marcel Jipp, Maximilian Ferle, Angela Rübeling, Tabea A. Oswald, Alf Honigmann, Andreas Janshoff

**Affiliations:** University of Göttingen, Institute of Physical Chemistry, Tammannstr. 6, 37077 Göttingen, Germany; University of Göttingen, Institute of Organic and Biomolecular Chemistry, Tammannstr. 2, 37077 Göttingen, Germany; Max Planck Institute of Molecular Cell Biology and Genetics, Pfotenhauerstraße 108, 01307 Dresden, Germany

**Keywords:** cell sorting, differential interfacial tension hypothesis, contractility, adhesion, tight junctions

## Abstract

Cellular sorting and pattern formation are crucial for many biological processes such as development, tissue regeneration, and cancer progression. Prominent physical driving forces for cellular sorting are differential adhesion and contractility. Here, we studied the segregation of epithelial co-cultures containing highly contractile, ZO1/2-depleted MDCKII cells (dKD) and their wildtype (WT) counterparts using multiple quantitative, high-throughput methods to monitor their dynamical and mechanical properties. We observe a time-dependent segregation process, governed mainly by differential contractility on short (< 5 h) and differential adhesion on long (> 5 h) time scales, respectively. The overly contractile dKD cells exert strong lateral forces on their WT neighbors, thereby apically depleting their surface area. This is reflected in a six-fold difference in excess surface area between both cell types. The lateral forces lead to a four-to sixfold increase in tension at all junctions that are in contact with the contractile cells including the interface between heterotypic cell-cell contacts. Concomitantly, the tight junction-depleted, contractile cells exhibit weaker cell-cell adhesion and lower traction force. Drug-induced contractility reduction and partial calcium depletion delay the initial segregation but cease to change the final demixed state, rendering differential adhesion the dominant segregation force at longer time scales.

This well-controlled model system shows how cell sorting is accomplished through a complex interplay between differential adhesion and contractility and can be explained largely by generic physical driving forces.

**Significance Statement:** Fundamental biological processes, such as tissue morphogenesis during development, rely on the correct sorting of cells. Cellular sorting is governed by basic physical properties such as the adhesion between cells and their individual contractility. Here, we study the impact of these parameters in co-cultures consisting of epithelial wildtype cells and overly contractile, less adhesive tight junction-depleted ones. We find time-dependent segregation into clusters: differential contractility drives fast segregation on short-time scales, while differential adhesion dominates the final segregated state over longer times.

## Introduction

Cellular sorting and tissue separation are essential processes in embryogenesis and tissue development, studied across multiple species.(1, 2) Early work has shown that cells taken from different embryonic tissues and remixed together eventually segregate again.(3, 4) Sorting of cells in tissues can be governed by different biological and physical factors. Owing to our accumulated knowledge about cell-cell junctions and the cytoskeleton, a hypothesis for cellular demixing based on differential adhesion was proposed.(5, 6) To accommodate different biological scenarios, this first hypothesis was complemented by incorporating differential cell contractility.(7, 8) Adhesion- and contractility-induced tensions act antagonistically: Contractility induces cell rounding to minimize the contact zone, whereas adhesion enlarges the cell-cell contact region. The resulting surface tension of the tissue is the ratio of adhesion and contractility.(9) This view has been extended more recently by the addition of local contractile cues, for example in the anteroposterior compartment boundary in *Drosophila*.(10–12) Alternatively, active cell forces have been proposed to also participate in regulating cellular demixing in co-cultures.(13) However, it remains difficult to differentiate between the various factors that govern cell sorting. In recent years, many simulation-based studies characterized different physical driving forces of demixing, identifying many possible pathways to cellular segregation via differential physical cell properties.(9, 14–22) Such simulations have the idea in common that the overall free energy in a cell layer, as determined by parameters such as contractility and adhesion, needs to be minimized. However, fundamental experimental evidence remains scarce, only applicable in certain scenarios, and often correlative.

Recently, it has been shown that in tight junction-depleted epithelial cells (ZO1 and 2 knockdowns; abbreviated as dKD) two distinct cell populations emerge. Some cells experience impaired ROCK signaling and contract, taking on a rounded shape; pulling on their neighbors eventually results in laterally elongated cells coexisting with the contracted cells.(23, 24) In initial experiments,(23) the stretched cell population was successfully replaced by less contractile wildtype (WT) cells, substantially increasing the mismatch in mechanical properties. In this co-culture, overly contractile dKD cells inhibited layer fluidity and migration through jamming. However, the driving forces for segregation in such a co-culture remain to be elucidated.

Here, we now address this question by studying co-cultures of dKD and WT cells using high-throughput/content (de-) mixing experiments in combination with quantitative mechanical single cell measurements. We focus on the quantification of cellular viscoelasticity, contractility, and cellcell adhesion to shine a light on the emergence and persistence of segregated cell monolayers. We found that a time-dependent demixing process in these co-cultured monolayers is governed by differential contractility on short time scales (within the first five hours), while on longer time scales (> 5 h) differential adhesion prevails. Such separation of demixing time scales has not been observed before. In addition, we show that the overly contractile dKD cells stretch out their WT neighbors and apically deplete their excess surface area, with a six-fold difference between the cell types. The dKD contractility leads to an about four-to six-fold increase in tension at all junctions in contact with these cells including the interface between heterotypic cell contacts. Additionally, the ZO1/2 depleted, highly contractile dKD cells exhibit weaker cell-cell adhesion and smaller traction forces. Taken together, our experimental results indicate that differential contractility prevails in the beginning to segregate the cell types, while with elapsed time differential adhesion becomes more and more important for demixing. Our results are consistent with Brodland’s differential interfacial tension model, which incorporates both contributions, but here leads to cell sorting in a time-shifted manner.(8)

## Results

### Demixing of co-cultured, highly contractile dKD, and compliant WT cells

First, live cell (de-) mixing experiments were recorded directly after thorough mixing and seeding using phase contrast and fluorescence microscopy (Figure 1A). We used WT cells with a GFP tag, (named WT-GFP from here on, see methods section) to distinguish them from dKD cells. Cell segmentation and neighbor analysis using both the fluorescence signal and phase contrast images allowed for the automatic assignment of cells as WT or dKD. This enabled us to quantify how much the cells mixed randomly or demixed into clusters, also called segregation. Therefore, we defined a segregation index *SI* as the number of homotypic neighbors divided by the number of all neighbors.

**Figure 1.**
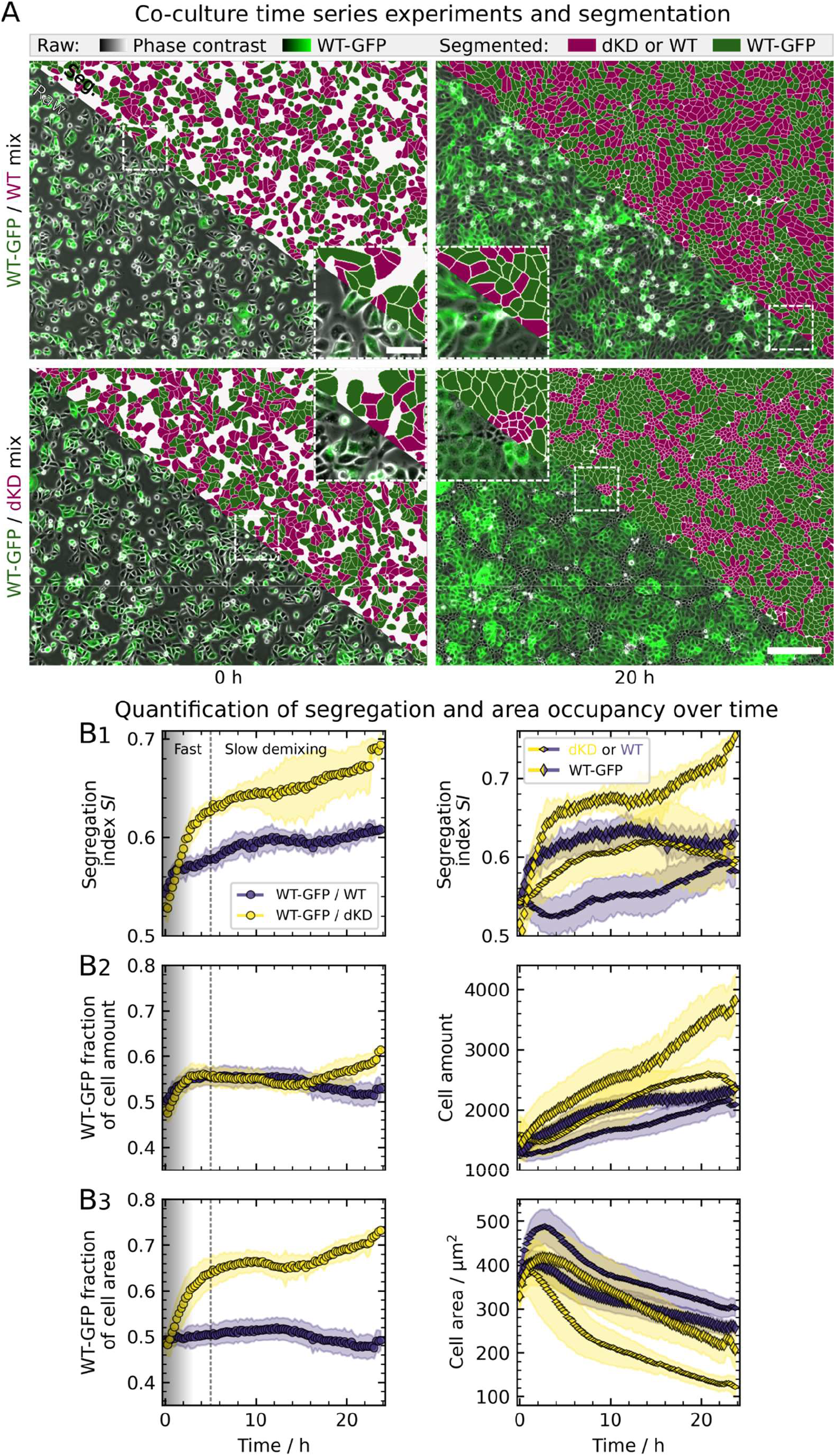
Demixing behavior of dKD and WT cell co-cultures at an initial mixing ratio of 50:50. A) Example overlay of phase contrast (grayscale) and fluorescence (green: WT-GFP cells) channels with corresponding segmentations (green: WT-GFP, magenta: dKD cells in WT-GFP/dKD mix or WT in WT-GFP/WT control, respectively). Samples were imaged immediately after seeding and mounting on the microscope (0 h). Scale bars: 200 μm and 50 μm (zoom-in). B) Demixing, cell amount, and area occupancy quantification. The vertical dashed line at 5 h indicates two distinct demixing time scales. The shade in the first 3 h indicates subconfluence. The yellow color represents the segmentation analysis of the WT-GFP/dKD mixture, while the purple symbols refer to the analysis of the control sample WT-GFP/WT. B1) The segregation index *SI*, defined as the average ratio of homotypic and all cell neighbors, quantifies the demixing degree. The *SI* is shown averaged over both cell types (left) as well as separately for each cell type (right). B2) Left: Relative cell amount, calculated as the ratio of the number of WT-GFP cells and the total cell amount. Right: Total cell amounts of each cell type. B3) Left: WT-GFP fraction of the overall cell area, calculated as the ratio of the WT-GFP area and the total cell area, indicating contractility discrepancies between the cell types. Right: Mean cell area of each cell type. Corresponding zoom-ins of the first 5 h are shown in Figure S1 and distributions of the individual cell areas are depicted in Figure S2. Mean values and standard deviations are shown. 12 separate regions from 6 culture dishes, acquired on three separate days (two regions per dish, two dishes per day), were measured and are shown per co-culture mix.

In the case of completely random cell distribution, an average segregation index of 0.5 would be expected. However, this parameter is also impacted by natural, local processes such as cell division. To account for these deviations from randomness, we performed control experiments using a pseudo co-culture consisting of WT-GFP cells and unmodified WT cells.

After detaching the cells and mixing the suspensions thoroughly before seeding, initially, both the WT-GFP/WT control as well as the WT-GFP/dKD mixture showed a segregation index close to 0.5 (Figure 1B1). The slight shift to higher values was likely already introduced upon initial seeding when most cells were still sedimenting, while others were already attached. Within the first hour, both co-cultures initially demixed from about 0.52 to 0.57 (Figure 1B1, see Figure S1 for zoomin). After this annealing time only the WT-GFP/dKD mixture segregated further, as expected. The SI increased to about 0.63 within the first 5 h, whereas the control remained at 0.57. After this fast initial demixing, both co-cultures segregated further at a similar rate to reach values close to 0.7 for the WT-GFP/dKD and 0.6 for the WT-GFP/WT cells.

In co-culture with dKD, WT cells are sorted into large, preferentially roundish clusters (Figure 1A) with a higher average SI than their dKD counterparts (Figure 1B1, on the right). The dKD cells, with a lower SI, were arranged in elongated, string-like clusters around the WT domains. This is expected from the differential interface tension model predicting the initial formation of chains of cells (here dKD) that in later stages anneal and coalesce.(8) If these later stages of complete segregation are reached depends on the difference in interfacial tension. In contrast, the WT-GFP/WT control showed an inconspicuous and less defined layer morphology. The *SI* of the labeled WT cells was generally higher than that of the unlabeled ones. However, this SI-difference vanished over time in the WT-GFP/WT co-culture, whereas in the WT-GFP/dKD mixture it even increased. Accordingly, WT-GFP/dKD co-cultures exhibited a sorting behavior into distinct clusters, different from homotypic monolayers.

In Figure S3 we show the detailed shape analysis and derived statistics of dKD clusters formed in confluent monolayers of WT and dKD cells (50:50) as a function of time compared with control samples composed of WT-GFP/WT cells (50:50). Initially (< 6 h), a larger number of clusters are formed compared with the WT-only sample, but they coalesce with time (> 6 h). This merging of clusters is attributed to the minimization of line tension generated by the contractility of dKD cells.(8) The aspect ratio follows this trend as it first rises and eventually decreases again.

As a control/normalization parameter for the *SI*, we next examined the cell amount of both cell types in each co-culture, because a difference in the relative cell amount could influence the *SI* as well. However, we observed no difference in the relative cell amount (WT-GFP fraction of the cell amount in Figure 1B2) between the WT-GFP/WT control and the WT-GFP/dKD cells. Interestingly, the total cell amount differed, with overall higher proliferation rates and larger cell amounts in the WT-GFP/dKD mixture. After a short delay in dKD cell proliferation, the dKD increased at a similar rate as the WT-GFP amount from about 3 h until 15 h after seeding. Importantly, the resulting small difference in the cell amount between the cell types was present in both the WT-GFP/dKD co-culture and WT-GFP /WT control (Figure 1B2), possibly slightly biasing the *SI* of both to larger values. After 15 h, dKD cells started to extrude apically out of the layer, offsetting proliferation and thereby stalling the cell amount. In the WT-GFP/WT mixture, the WT-GFP also showed slightly more proliferation until 15 h after seeding, which then leveled off.

Next, to examine the cell contractility discrepancy of these cell lines, which was described previously(23, 24), we first quantified the labeled WT fraction of the cell area (Figure 1B3). If there were no discrepancies in contractility in the co-culture, this parameter would be expected to be 0.5 because each cell type would occupy 50% of the covered area. Indeed, this was the case for the WT-GFP/WT control. In contrast, however, the WT-GFP/dKD co-cultures showed a strong increase in the WT area fraction within the first 5 h, precisely correlating with the *SI* increase (see Figure S1 (zoom-in)). This highlights a great differential contractility with highly contractile dKD cells occupying smaller areas and stretched WT cells covering more space on the culture dish. At the same time, as described before, the relative cell amount stayed constant, confirming that the larger area coverage of WT cells is due to lateral extension provoked by contractile dKD cells and not a consequence of an increased amount of WT cells. Notably, this effect only develops over time due to collective cell-cell interactions because the WT-GFP/dKD mixture also starts at *SI* = 0.5. However, the contractile discrepancy is generally underestimated here. This is because the phase contrast channel was used for analysis (the fluorescence was only used to assign the cell type, see methods section) but the lateral stretching of bordering WT cells by dKD neighbors can be best observed in the WT-GFP specific fluorescence channel (white arrows in Figure 2B). This is because the WT cell body extension, even overlapping above dKD cells, is specifically seen in the GFP channel (Figure 2B) while in phase contrast, the overlapping WT and dKD cell bodies cannot be distinguished well (Figure 1A).

**Figure 2.**
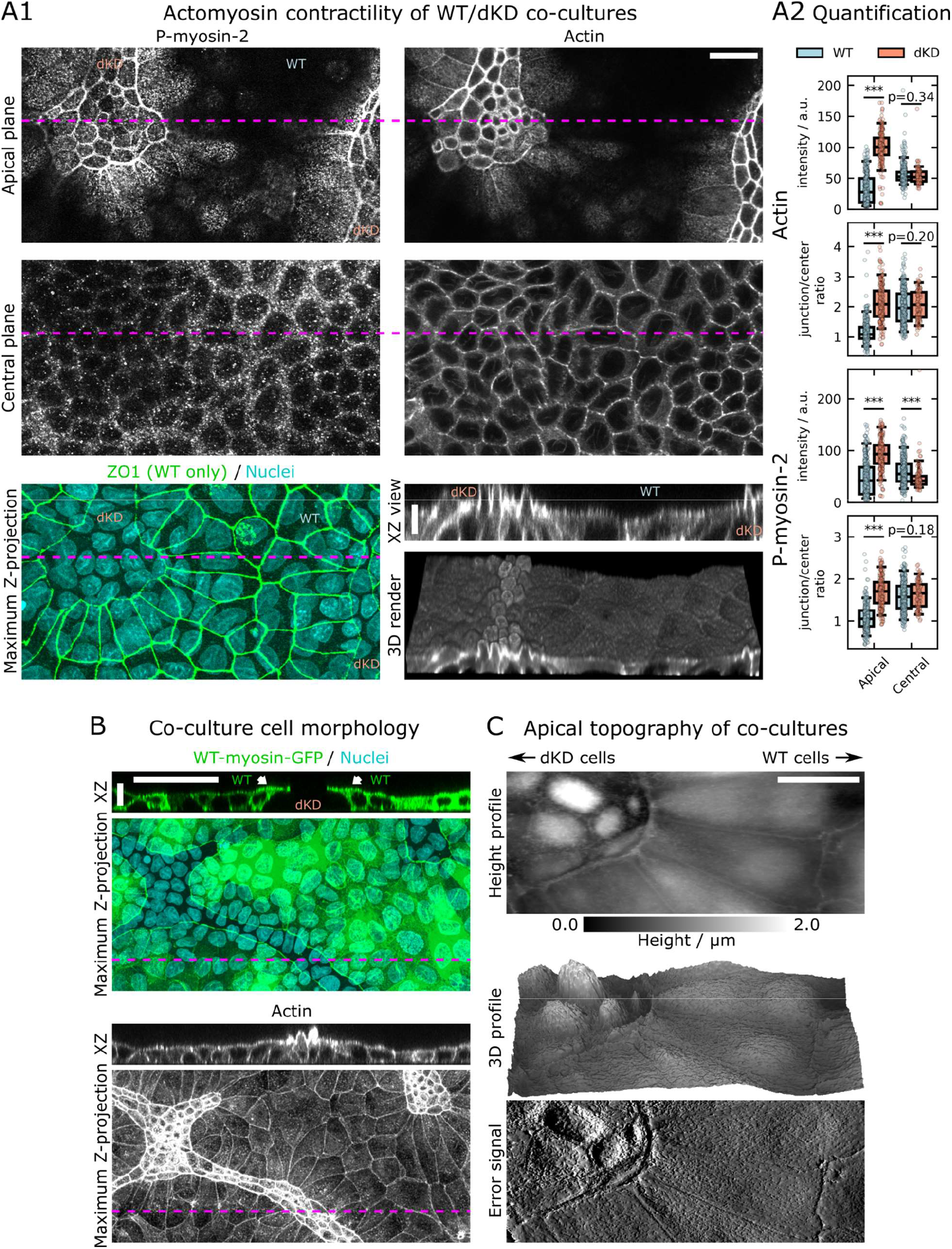
Differential actomyosin contractility of WT-GFP/dKD co-cultures. A1) Representative WT-GFP/dKD co-cultures co-stained for phosphorylated myosin (P-myosin-2 antibody), actin (phalloidin), nuclei (DAPI, cyan), and ZO1 (ZO1 antibody, green). ZO1 was used to distinguish ZO1/2 dKD from WT cells. Magenta lines indicate the location of the XZ view. XY scale bar: 20 μm, Z: 5 μm. A2) Quantification of fluorescence intensities from A1 including data from two large images. Boxes show the median and the upper and lower quartiles, whiskers indicate the 5th and 95th percentile, while data points represent individual cells. Junction to center ratio refers to fluorescence intensity ratios obtained by a segmentation process detailed in the SI. B) WT-GFP/dKD co-culture co-stained for actin (phalloidin) and nuclei (DAPI, cyan). The green channel was used to identify the WT-GFP cells and to examine their morphology in 3D. XY scale bar: 50 μm, Z: 10 μm. C) Apical topography of WT-GFP/dKD co-cultures obtained by AFM imaging. Height profile, the corresponding 3D topography map which was up-scaled vertically by 50%, and the error signal (deflection image). Scale bar: 20 μm. Cells in A were fixed after 28 h, and in B and C after about 48 h of growth.

Generally, the cell area of both WT and dKD cells in co-culture or control samples decreases over time due to the compaction and jamming of cells in the confluent state (Figure 1B3 (right panel)). After 18-20 h jamming has finished and due to the strong contractile forces, extrusion of primarily dKD cells leads to an overall loss of cells. After approximately 20 h the number of dKD cells decreases due to preferential exclusion from the cell monolayer, a consequence of apical constriction. Notably, extruded cells are sometimes not correctly captured by the segmentation algorithm, leading to an overall decrease of occupied area.

### Impact of proliferation on the segregation index

Besides contractility and adhesion, proliferation might also be an important factor to foster demixing by enlarging the clusters in the demixed state. Albeit we could not find an enlargement in dKD cluster size (see Figure S3) we conducted experiments in the presence of mitomycin C which effectively suppresses proliferation in the separation process (Figure S4). Switching off proliferation essentially had no impact on initial cell sorting (Figure S5). However, cell sorting at later time points is slowed, suggesting missing dKD cells that were removed from the cell layer due to apical constriction in the untreated WT-GFP/dKD (Figure S4) as also discussed above.

### Differential actomyosin contractility and 3D cell morphology of WT-GFP/dKD co-cultures

To further study the differential contractility of WT-GFP/dKD co-cultures on a molecular and cell morphological level, we applied confocal fluorescence microscopy and AFM imaging (Figure 2).

In previous work(23), we observed a strong actomyosin upregulation at the apical-lateral cell periphery of dKD cells (Figure 2A). Particularly activated (phosphorylated) myosin accumulated at the apical cell-cell junctions. A thick perijunctional actomyosin ring was formed, constricting the ZO1/2 depleted cells apically. Conversely, the WT cells did not show any upregulation of phosphorylated myosin-2 or of the actin cytoskeleton. To conserve the cellular volume, dKD cells were forced to bulge out apically. Since all dKD cells were still connected to their neighbors, adjacent WT cells were stretched out and flattened by the apical pull of the dKD cells. Strikingly, WT cells at the WT-GFP/dKD interface were partially pulled across their direct dKD neighbors towards the center of the dKD cluster (Figure 2B, white arrows). Note that this lateral pulling translocates certain cell components, such as ZO1 or myosin in Figures 2A1 and B, relative to the nucleus. Both for actin and p-myosin it was found that dKD cells display a larger junction to center ratio of fluorescence intensity compared with WT cells at the apical plane (Figure 2A2). The lateral elongation of WT and apical contraction of dKD cells was confirmed by AFM imaging (Figure 2C). Interestingly, the bordering junctions at the interface between a WT and dKD cluster are particularly pronounced on the apical side (Figure 2C). This was reflected by increased myosin accumulation in this region (Figure 2B), which overall highlights the mechanical discrepancy between cell types.

### Differential mechanics of dKD and WT cells in co-culture

To directly quantify the mechanical consequences of the described contractile, molecular, and morphological disparities in WT-GFP/dKD co-cultures, we examined their mechanical phenotypes by AFM indentation-relaxation, traction force microscopy as well as laser ablation (Figure 3).

**Figure 3.**
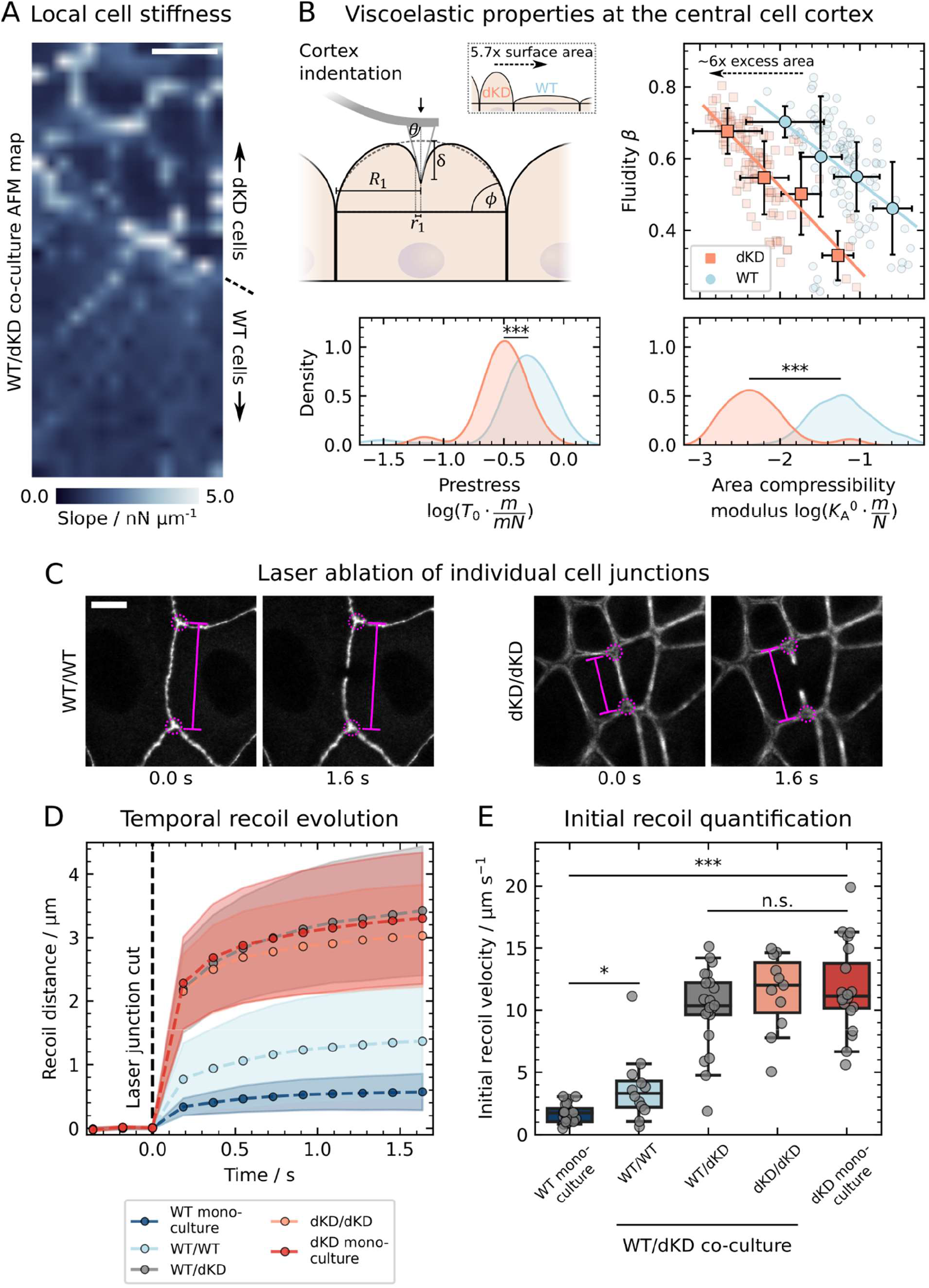
Differential mechanical properties of dKD and WT cells in co-culture. A) AFM map showing the slope of the force curve during contact, which locally reflects the apparent mechanical stiffness. Scale bar: 10 μm. B) Site-specific viscoelastic properties of the central cell cortex in proximity to the WT-GFP/dKD interface. The cortex indentation geometry considered in the so-called Evans model includes the contact angle *ϕ* and base radius *R*_1_ of the spherical cell cap, the indentation depth *δ*, and the contact radius *r*_1_. Importantly, dKD cells had a pronounced cap with larger *ϕ* and smaller *R*_1_ than WT cells (vide supra), yielding a 5.7-fold surface area difference. Upon fitting, the fluidity *β* was plotted against the decade logarithm of the scaling factor representing the area compressibility modulus *K*_A_^0^, and histograms for the latter and the prestress T0 are shown. Small transparent data points represent individual indentations. Large symbols and error bars are binned means and standard deviations. Lines indicate linear fits (in log space) of the binned means. C) Laser ablation examples of individual cell junctions. In WT cells, ZO1 in the tight junctions was stained, and in dKD cells, myosin was stained. D) Tensile junction properties were obtained by tracking the distance (magenta lines) between two opposing junction vertices (magenta circles) upon recoil. Temporal means and standard deviations are shown. E) The initial recoil velocity was calculated between the last point before (0.00 s) and the first one after ablation (0.18 s). The boxes show the median and the upper and lower quartiles. Whiskers indicate the 5th and 95th percentile, while data points represent a single cut of one junction. Scale bar: 10 μm. All measurements were repeated in at least three independent experiments and performed on multiple WT-GFP/dKD clusters.

First, we acquired AFM indentation maps (Figure 3A) and examined the apparent local stiffness, which is reflected in the slope of the force-distance curve. Here, we observed a similar picture as in pure dKD monolayers,(23) dKD cells were softer at the central cortex and extremely stiff at the perijunctional actomyosin ring (vide supra). In contrast, neighboring WT cells showed only slightly pronounced cell boundaries but an increased stiffness at the center in comparison with dKD neighbors.

To further characterize this stiffness difference at the center of the two cell types, we performed site-specific indentation experiments followed by force relaxation and applied a tailor-made viscoelastic fitting model as described in several recent studies.(23, 25, 26) In brief, this model fits the stress relaxation of the composite viscoelastic shell upon indentation according to a power law of the area compressibility modulus assuming constant volume during the experiment (see SI for a detailed description of the mechanical model). Importantly, the cell geometry (area and angle of the apical cap), which differs tremendously between both cell lines (vide supra), can be adjusted in this model (Figure 3B). We limited the indentation depth to about 1 μm to minimize the impact of the nucleus on the cellular response to deformation (see Figure S6 for geometric reasoning). Three parameters are obtained: the prestress *T*_0_ corresponding to the actomyosin cortex tension, the apparent area compressibility modulus (scaling factor) *K*_A_^0^, which besides the shell’s stiffness also mirrors the excess cell surface area, and the fluidity *β* representing the viscous behavior (energy dissipation) of the cortex. *β* = 1 corresponds to a Newtonian fluid, whereas *β* = 0 refers to an elastic solid.

Because *K*_A_^0^ was previously found not to be independent of but to scale with the fluidity *β*,(26) we plotted *β* against *K*_A_^0^ (Figure 3B). Interestingly, we found that the fluidity was not significantly different between WT and dKD cells (p = 0.53, with the same median of 0.6). However, we found a shift to larger *K*_A_^0^ values for WT cells compared with their dKD neighbors in co-culture. This increase can be attributed to the removal of excess surface area Aex compared with the geometrical surface area *A*_0_ of the cells via 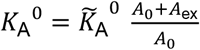.(27)

The picture which therefore emerges suggests that surface area is sacrificed to mitigate the external stress from adjacent dKD cells. This occurs at the expense of cell stiffening but preservation of fluidity. On one hand, we observed an unchanged fluidity and only a relatively small difference in prestress within the range of the standard deviation (0.49 ± 0.22 mN m^-1^ for WT cells compared with 0.31 ± 0.14 mN m^-1^ for dKD (median ± s.d.)). On the other hand, WT cells exhibited a substantially larger scaling factor *K*_A_^0^, with an increase of more than one order of magnitude (0.061 ± 0.084 mN m^-1^ for WT compared with 0.004 ± 0.016 N m^-1^ for neighboring dKD cells (median ± s.d.)). From *K*_A_^0^ we were able to estimate that about six-fold as much excess surface area was stored in dKD as in WT cells. This fits the theoretical 5.7-fold surface area difference between the different cap geometries, indicating that the apical surface material is conserved upon stretching, i.e., dKD cells contract laterally and store the membrane/cortex material apically, while WT cells sacrifice apical excess area to cope with the external areal strain. Although a tremendous amount of excess surface material is sacrificed by WT cells, the mechanics of the cortex is largely unaffected, with small differences in prestress. In agreement, we also did not observe an obvious change in the actin signal at the central cortex in Figure 2A (*vide supra*). In consequence, the observed lateral contraction of the dKD cells did not originate simply from cortex mechanics but most likely from the perijunctional actomyosin ring (Figure 2A).

To confirm this assumption and to characterize how the differential contractility translates into interfacial tension in the layer, we specifically examined the junctional tension using laser ablation, severing the cell junctions (Figure 3C-E). Particularly, we addressed the tensile properties of the cell junction between a WT and another WT cell, dKD/dKD junctions as well as the WT-GFP/dKD interface. In addition, we compared the new mechanical equilibrium in co-cultures with the tension of the junction in WT and dKD monocultures. For this purpose, we analyzed the recoil dynamics of the opposite apex nodes of the ablated compound over time (Figure 3D) and recorded the initial recoil velocity (Figure 3E).

We found a significant, four-to six-fold increase in recoil velocity for all junctions bordering a contractile dKD cell (10-12 μm s^-1^, compared with 2-3 μm s^-1^ without any direct contact with a dKD cell). dKD/dKD junctions in co-culture were comparable with dKD mono-cultures (*p* = 0.83). Interestingly, the WT-GFP/dKD interface had slightly smaller recoil velocities than dKD/dKD junctions (*p* = 0.29), while WT/WT junctions displayed slightly but significantly higher velocities than WT mono-cultures. This highlights the establishment of a new mechanical equilibrium in cocultures based on a tug-of-war between highly contractile dKD cells and compliant WT neighbors; in co-cultures, tension from dKD cells is accommodated by WT cells, while in dKD mono-cultures all cells exhibit increased tension. Forces are transmitted over long distances within WT clusters to distribute the load and the whole cell monolayer is under extensile tension as all cut junctions display the same recoil direction (retraction towards the junction knots). dKD cells pull at their neighbors thereby generating elevated tension at the junctions. The whole cell monolayer is under extensile tension, like a fluid wetting a surface. Overall, the data shows that the increased contractility of dKD cells translates into increased junctional tension of all direct neighbors in the layer, while compliant WT cells serve to buffer the lateral stress.

A further indication that the observed segregation is based on interfacial tension, was another set of experiments in which we varied the mixing ratio between dKD and WT cells before seeding (Figure S2). We observed that the pattern of elongated dKD cell stripes which surrounded the predominately roundish WT clusters persisted, independent of the mixing ratio. This is indicative of interfacial energy minimization in accordance with the tension-based sorting hypothesis and is in contrast to demixing driven by active forces as reported recently.(13)

Traction force microscopy was carried out to measure the impact of the cytoskeletal remodeling in response to ZO1/2-depletion on the cell-substrate interaction. We found that the traction forces (per unit area) exerted by confluent WT cells were more than twice as high (84.0 ± 19.5 Pa (mean ± s.d.), *n* = 7 monolayers) as those observed for confluent dKD cells (38.5 ± 5.2 Pa (mean ± s.d.), *n* = 8 monolayers). This suggests that the remodeling of the actin cytoskeleton also involves the basal side, creating a strong imbalance between the apical and basal side, and goes hand in hand with the reduced cell-cell adhesion measured between pairs of dKD cells (*vide infra*).Likewise, larger adhesion forces are required to balance the strong forces exerted by WT cells on the substrate.

### Differential cell-cell adhesion of WT and dKD cells

While the increased contractility of dKD cells is well documented and could induce segregation via energy minimization, changes in intercellular adhesion might also be expected due to the loss of the adhesion-mediating junctional ZO proteins.

To quantify cell-cell adhesion, we performed AFM experiments with one cell attached to the AFM cantilever serving as the probe and the other one adhered to the Petri dish. The two cells were brought into conformal contact and separated after a short dwell time in contact (Figure 4A). The separation forces between the two cells are not obtained under equilibrium conditions and are therefore referred to as the dynamic adhesion strength. We found decreased adhesion forces for all dKD cells (two dKD cells as well as a dKD adhering to a WT-GFP cell) as shown in Figure 4B. As a control, we also compared WT cells and GFP-tagged WT cells. While the WT-GFP cells displayed slightly lower adhesion forces than pure WT cells, they are still consistently more adhesive than dKD cells (p < 0.001 compared with WT-GFP/dKD and *p* < 0.01 with dKD/dKD). Interestingly, the adhesion between two cells was always dominated by the respective weaker binding partner, i.e., the dKD cells, indicative of largely immobile receptor-ligand pairs. Accordingly, in WT-GFP/dKD co-cultures differential adhesion and contractility together determined the differential interfacial tension during cell segregation.

**Figure 4.**
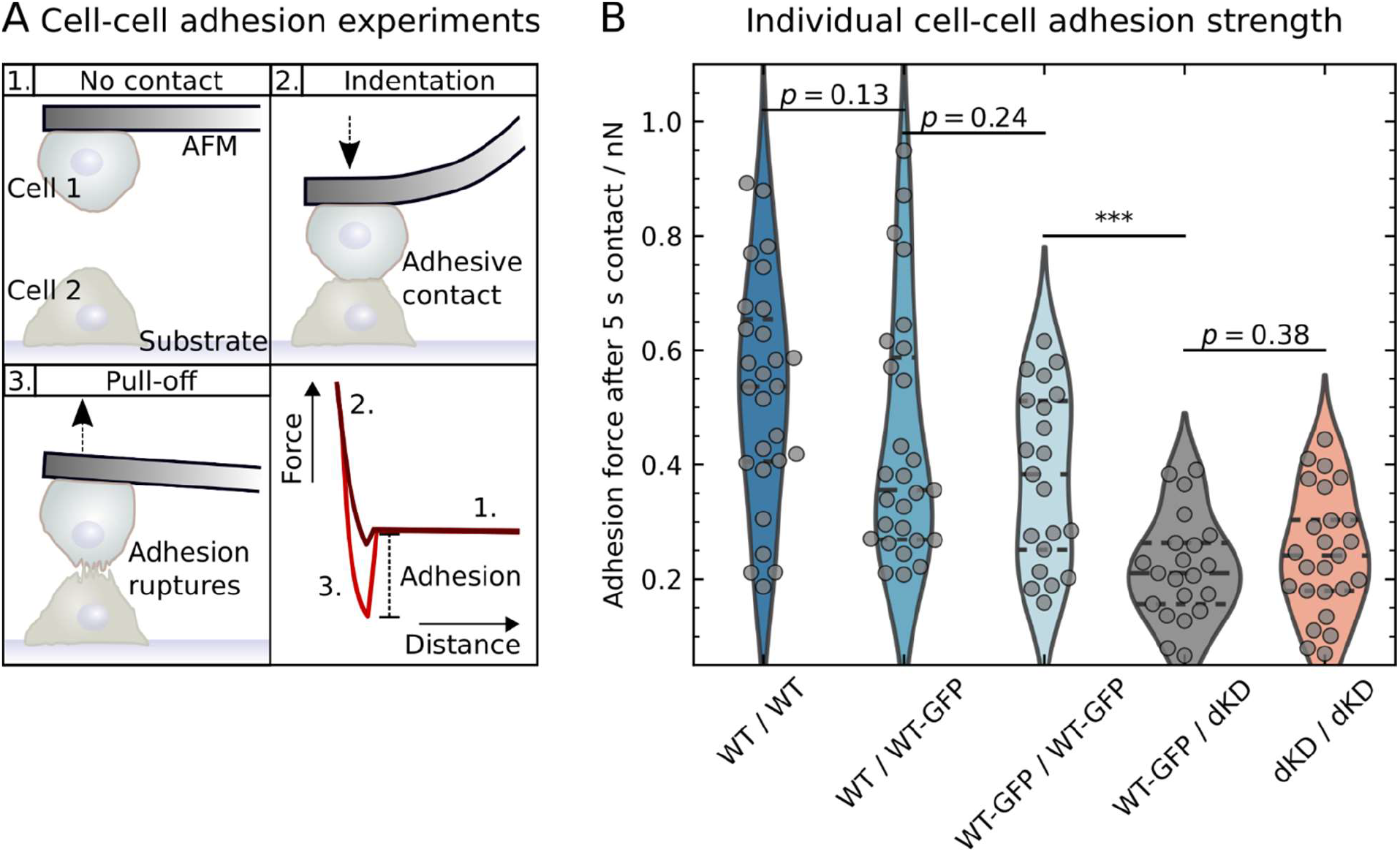
Differential intercellular adhesion of WT and dKD cells. A) AFM-based adhesion measurements: 1. Before or after an experiment, one cell is connected to the cantilever and one adheres to the culture dish substrate, without contact with each other. 2. Adhesive contact between cells is established at 2 nN for 5 s. 3. As the cantilever is retracted, the cells are pulled apart and bonds rupture. Schematic retraction curves depict small (dark red) and large (light red) adhesion forces. B) These adhesion forces are compared between different important cell combinations. Violins represent kernel density estimation with horizontal, dashed lines showing the quartiles and median. Violins are scaled to have the same area. Single data points represent individual adhesion peak forces. Three consecutive indentation/retraction cycles were performed for each cell pair. For each combination, at least 4 individual cell-cantilever probes, and at least 8 cells on the substrate were measured, with experiments repeated on at least 4 days.

The loss of ZO proteins perturbs tight junction-associated signaling by redistributing ROCK to the adherens junction,(24) where it leads to apical constriction in dKD cells that coexist with outstretched WT cells to maintain force balance and avoid bending of the 2D monolayer away from the surface. Myosin-2 is also redistributed and predominately found in the apical region (Figure 2A1) where it fosters constriction, while concomitantly adhesion to the substrate is diminished.

### Time-scale dependency: Contractility drives early, adhesion final sorting

While we established that there is differential contractility and differential adhesion in WT-GFP/dKD co-cultures, it remains unclear, which one dominates over the other. Therefore, we performed the demixing experiments shown in Figure 1 again, however, this time in the presence of the Rho kinase (ROCK) inhibitor Y27632, to reduce cell contractility, switching off one of the contributions to demixing (Figure 5, Figure S7-8). Y27632 mainly affects the actomyosin contractility of cells, while the difference in cell-cell adhesion should remain the same. This should restore the apical conformity lost due to the increased ROCK activity in response to ZO1/2 depletion. We also impaired all cell-cell contacts by reducing the calcium content of the medium to induce a similar effect (Figure S9). Here, the effective differential contractility was reduced concomitantly (Figure S9B, bottom). Importantly, upon completely abolishing both contractility and adhesion (at 0 mM Ca^2+^), no sorting was possible neither in WT-GFP/WT nor in WT-GFP/dKD cultures, leading to *SI* levels even lower than WT-GFP/WT controls in normal medium (Figure S9B, top).

**Figure 5.**
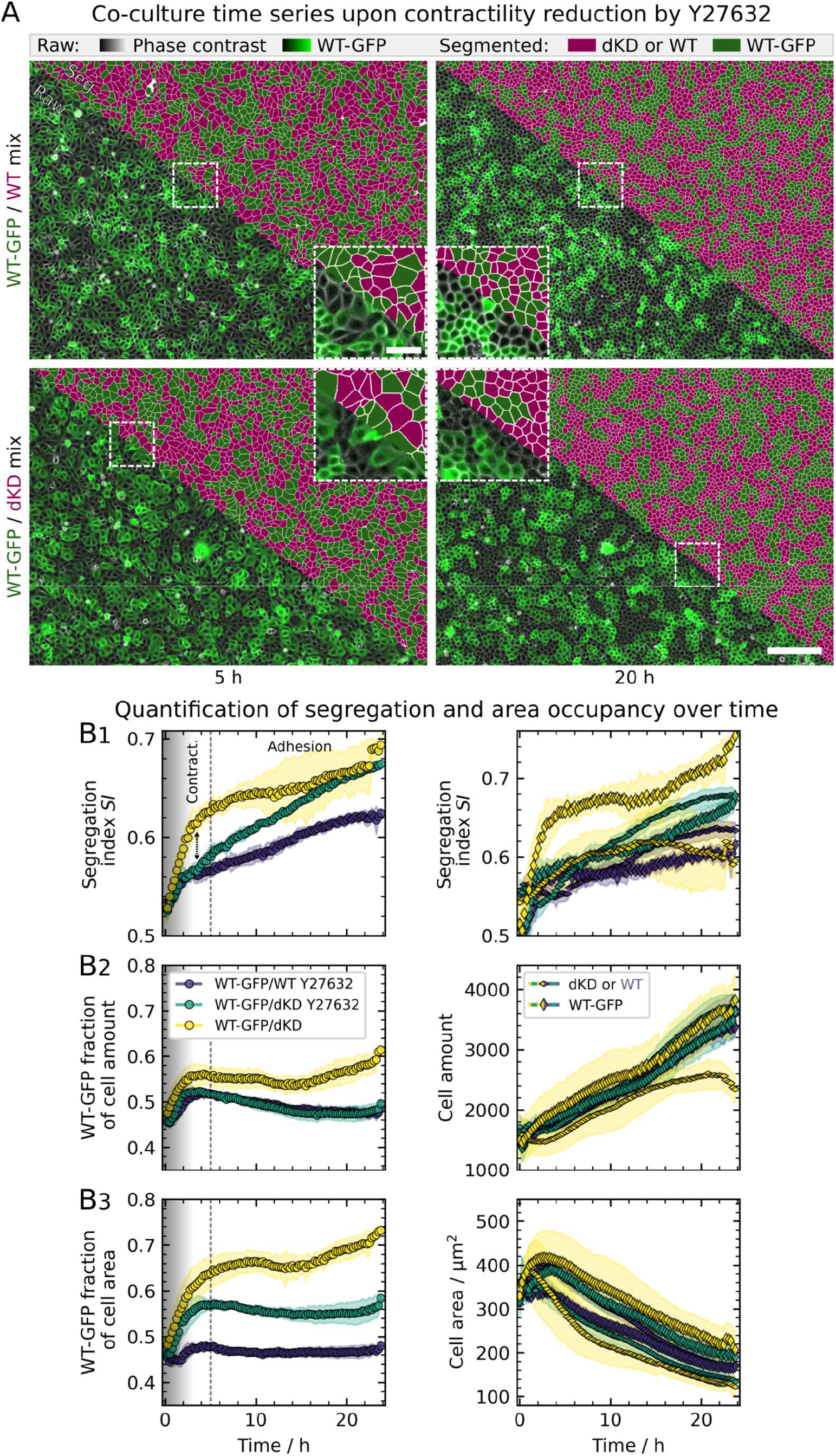
Contractility drives early, adhesion final sorting. Demixing behavior of highly contractile dKD and wildtype cell co-cultures at an initial mixing ratio of 50:50, treated with 50 μM Y27632. Experiments and Figure panels are set up analogously to Figure 1, and, for comparison, untreated WT-GFP/dKD from Figure 1 were included. A) Example overlay of phase contrast (grayscale) and fluorescence (green: WT-GFP cells) channels with corresponding segmentations (green: WT-GFP, magenta: dKD cells in WT-GFP/dKD mix or WT in WT-GFP/WT control, respectively). Samples were imaged immediately after seeding and mounting on the microscope (0 h). Images from the start of the experiment (t = 0 h) can be found in Figure S7A. Scale bars: 200 μm and 50 μm (zoom-in). B) Demixing, cell amount, and area occupancy quantification. The vertical dashed line at 5 h indicates two distinct demixing time scales, thought to be determined by contractility and adhesion. The shade in the first 3 h indicates subconfluence. Yellow symbols represent data from untreated WT-GFP/dKD mixtures, purple symbols refer to with ROCK inhibitor Y27632 treated WT-GFP/WT control and green symbols refer to WT-GFP/dKD cell mixtures exposed to Y27632. B1) The segregation index SI, defined as the average ratio of homotypic and all cell neighbors, quantifies the demixing degree. The *SI* is shown averaged over both cell types (left) as well as separately for each cell type (right). B2) Left: Relative cell amount, calculated as the ratio of the number of WT-GFP cells and the total cell amount. Right: Total cell amounts of each cell type. B3) Left: WT-GFP fraction of the overall cell area, calculated as the ratio of the WT-GFP area and the total cell area, indicating contractility discrepancies between the cell types. Right: Mean cell area of each cell type. Corresponding zoom-ins of the first 5 h are shown in Figure S7B, a comparison of untreated and Y27632-treated WT-GFP/WT cultures as well as later experiment times (24 h - 35 h) can be found in Figure S8, and distributions of the individual cell areas are depicted in Figure S2. Velocity and persistence analyses can be found in Figure S10. Mean values and standard deviations are shown. 6 separate regions from 3 culture dishes (two per dish), acquired on separate days, were measured and are shown per co-culture mix.

Upon first visual inspection after ROCK inhibition (Figure 5A), at early time stages, no difference was discernible between the WT-GFP/dKD mixture and the WT-GFP/WT control. Only at later times, stronger demixing was observed in the WT-GFP/dKD co-culture as mirrored in the segregation index (Figure 5B1). Here, we plotted the untreated WT-GFP/dKD mixture from Figure 1 to serve as a reference, together with contractility-inhibited WT-GFP/dKD and WT-GFP/WT cocultures. While the WT-GFP/WT control did not change its segregation behavior upon Y27632 administration, the very fast, early segregation of WT-GFP/dKD co-cultures (< 5 h) was substantially diminished. Instead of this fast initial behavior, segregation of the WT-GFP/dKD mixture was slowed down. Nevertheless, after about 15 h the contractility-inhibited WT-GFP/dKD mixture reached approximately the same *SI* of approximately 0.7 as the untreated counterpart. Accordingly, the upregulated contractility of dKD cells was critical for early segregation, while the adhesion differential was still able to induce cellular demixing upon longer time scales.

As a control parameter, we also inspected the ratios of cell area and amount (Figure 5B2-3) as in Figure 1. The WT fraction of the cell amount (Figure 5B2) again served to provide context for the *SI* values and relative area coverage. While the untreated WT-GFP/dKD cell amount ratio was slightly shifted towards more WT cells, both drug-treated co-cultures remained at a 0.5 ratio (Figure 5B2). Interestingly, the proliferation in the WT-GFP/WT control was increased by Y27632 to the same level present in treated and untreated WT-GFP/dKD (except for the dKD extrusion after 15 h) as shown in Figure 5B2 and Figure S10A, while the *SI* remained much lower. To further rule out that local clustering due to proliferation dominates the segregation, we investigated the relationship between the *SI* and the cell amount (Figure S10A) upon exposure to Y27632. The *SI* generally increased with increasing cell amounts but with a lower slope at higher cell amounts. However, while for both treated cultures and the WT-GFP/dKD mixture the proliferation rate was approximately constant over time, the scaling of the *SI* with the total cell amount was much different. At the same cell amount, the *SI* remained lower in the treated WT-GFP/WT control than in the untreated WT-GFP/dKD mixture. In addition, the difference in proliferation between the treated and untreated WT-GFP/WT samples did not translate into an increase in segregation. Note that the cell amount is essentially equivalent to cell density in our experiments because the size of the field of view was always the same.

To assess the cell contractility the area ratio once again served as a broad-scale readout (Figure 5B3). Here, we did observe the expected drop upon contractility inhibition for the WT-GFP/dKD mixture, while the WT-GFP/WT control was unaffected. Importantly, this drop in contractility remained over the whole duration of the experiments, confirming that the effect of the drug did not wear off over time. Moreover, since switching off proliferation by administration of mitomycin C (*vide supra*, Figure S4) neither changed cell sorting dynamics, we can safely conclude that first contractility and later adhesion dominate the segregation process.

To rule out that the drug acts on cell motility (e.g., due to effects on the focal adhesions on the substrate) influencing demixing, we quantified the velocity and persistence via cell tracking (Figure S10B). We investigated this, particularly for the first 5 h, where the impact of the drug on segregation is the strongest. If higher motility was a driving factor for random mixing, we would expect an increase in the motility parameters, particularly of the WT-GFP/dKD mixture upon drug treatment. However, this was not the case, but, to the contrary, the motility parameters even decreased slightly or remained the same (Figure S10B). The WT-GFP/WT control showed a slight drop in both parameters, while its (de-) mixing behavior was largely unaffected. Accordingly, the drug provoked the delay in WT-GFP/dKD sorting not by affecting motility but indeed via inhibiting cellular contractility.

Additionally, we also confirmed that inhibition of ROCK does not significantly alter cell-cell adhesion. For this purpose, we cultured WT-GFP cells on an AFM cantilever facing with the apical part to the opposing monolayer on the Petri dish (μ-dish low; ibidi), and measured separation forces in the presence and absence of Y27632 (Figure S11). We found that adhesion between WT-GFP and dKD cells is only slightly reduced by the ROCK inhibitor (330 ± 130 pN down to 300 ± 120 pN with Y27632 (mean ± s.d.)) at 5 s contact time between the cells. This is, however, also true for the separation force measured between two WT-GFP cells (435 ± 350 pN to 370 ± 200 pN with Y27632 (mean ± s.d.)), which means that the overall impact of Y27632 on cellcell adhesion is small and the gradient of adhesion strength between the two cell types remains unchanged in the presence of ROCK inhibitor.

A similar effect was obtained with reduced calcium concentration in the culture medium, slowing down the contractility-based cell sorting (Figure S9, 0.07 mM Ca^2+^) without inhibiting demixing, i.e., the same final *SI* is reached after > 20 h. However, further withdrawal of calcium from the culture medium completely abolished both contractility and adhesion-based cell sorting. As expected, the two cell types do not display any segregation anymore.

Together, data from these experiments showed that co-cultures can display an intricate interplay of contractility and adhesion driving cell segregation on distinct time scales.

## Discussion

Our goal was to identify and scrutinize the driving forces for the demixing of co-cultures consisting of WT and dKD MDCKII cells displaying both different cell-cell adhesion due to the knock down of ZO1/2 and differential contractility due to actomyosin upregulation in the apical domain. Tight junction-associated signaling is pivotal for maintaining epithelial sheet morphology, integrity, and function.(28–32) In planar epithelial cell sheets the apical, contractile forces are typically balanced to avoid deformation and bending of the entire sheet. The Par polarity proteins Par-3, and Willin, a FERM-domain protein, are involved in this regulation by suppressing the junctional localization of ROCK through its phosphorylation by the protein kinase aPKC.(24, 28) This ensures uniformly shaped apical domains and balanced contractility. Loss of ZO1/2, however, perturbs Par-3 localization and therefore leads to apical constrictions and atypical apical morphology.(24) ZO proteins are required for epithelial polarization and it was shown previously that depletion displays unbalanced tensile strain from surrounding cells that could, however, be largely restored by inhibition of ROCK.(23, 24) It was therefore of great interest to examine the consequences of ZO depletion for cell sorting and layer morphology.

We found that the main driving forces for creating clusters of dKD cells coexisting with WT clusters are time-scale separated. On short time scales (within the first five hours) differential contractility prevails, while on longer times scales (>5 h) cell sorting is driven predominately by differential adhesion. Clusters of dKD cells are shaped by elongated chains of cells that shorten in later stages. This dynamic behavior is expected from the general rule that cell sorting occurs in three distinct steps that include the formation of elongated chains that shorten and smooth and finally, when the tension between the different cell types is large enough compared to the homotypic tensions, anneal into large round clusters and minimize the line tension.(8) To our knowledge, this is the first time that a separation of time scales in cellular segregation is described and attributed to distinct mechanical properties. Our data suggest that if differential contractility is abolished, differential adhesion alone is sufficient for cell sorting but considerably slower and less efficient.

The envisioned mechanism comprising adhesion- and contractility-based cell segregation is summarized in Figure 6. While in randomly mixed WT cultures adhesion between all cells is the same and they display similar contractility, in WT-GFP/dKD co-cultures the adhesion and contractility between the cell types are considerably different, inducing segregation into clusters. In particular, the difference in apical contractility between dKD and WT cells results in a substantial tension difference that initially favors rapid but partial sorting. The apical forces are balanced by neighboring cells, so that large, outstretched WT cells coexist with small, contractile dKD cells. The apical forces exerted by the dKD cells are balanced by WT neighbors via their higher traction forces on the substrate. This force balance prevents bending of the cell layer into the third dimension. Cell-substrate adhesion forces are stronger for WT cells than for dKD cells, as also suggested by phosphorylated myosin-2 staining (Figure 2). In response to the contraction of adjacent dKD neighbors, WT cells are stretched and therefore sacrifice a large amount of excess surface area to prevent lysis. If contractility is balanced again by inhibiting ROCK, segregation based on differential contractility is strongly delayed. The same was found for moderate calcium depletion slowing down initial cell sorting by decreasing the effective differential contractility (Figure S9B). It was shown that Y27632 restores normal apical area distribution in ZO1/2 depleted Eph4 cells.(24) The authors found an excess of contractility due to aberrant ROCK activation at the adherens junctions. However, even after switching off contractility, the remaining tension difference due to stronger WT-GFP/WT adhesion compared with dKD/dKD adhesion still induces the same amount of segregation as in untreated layers over a longer time, highlighting a redundant but time-dependent role of contractility and adhesion. Considering that adhesion complexes mature progressively over time,(33) whereas contractility is a property of individual cells, it is conceivable that differential contractility promotes sorting immediately while adhesion acts on longer time scales. Depletion of calcium to an extent that prevents cells from ablation led to full suppression of demixing as it switches off both cell-cell adhesion and contractility. Using only moderate calcium depletion we found that only the contractility-based cell sorting was affected, i.e., slowing down initial cell sorting (Figure S9, presumably via calcium’s role in promoting cell contraction (34, 35)). Proliferation plays only a role in later stages when jamming occurs and dKD cells are extruded from the cell layer due to the strong apical forces (Figure S4).

**Figure 6.**
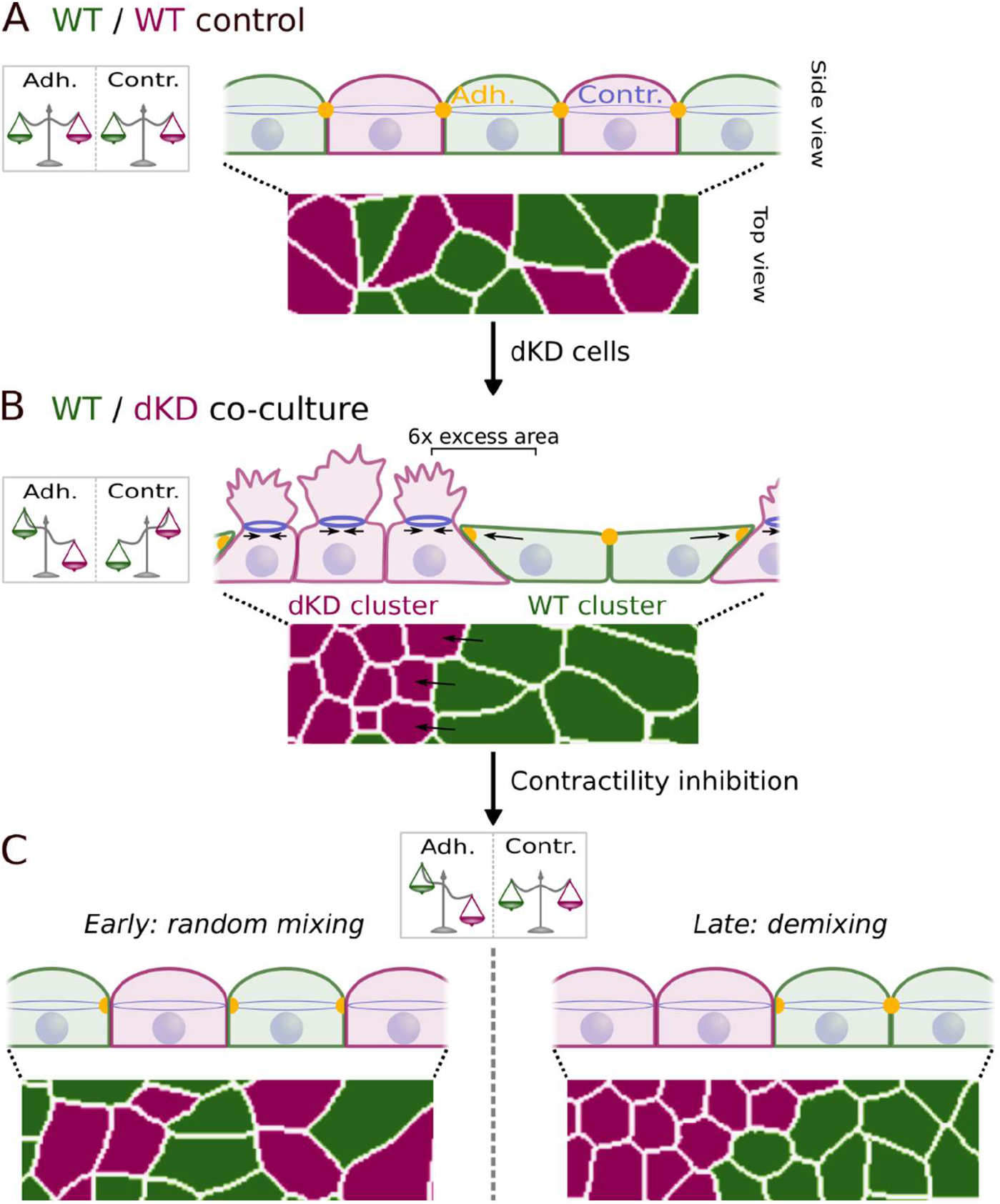
Proposed model of the interplay between adhesion and contractility in WT-GFP/dKD cell layers. A) In WT-GFP/WT control layers, adhesion is the same between all cells and they are equally contractile, hence, random mixing takes place. B) Adding dKD cells induces differences in both adhesion and contractility between the cell types. dKD cells lose some adhesive contact and contract excessively, yielding tremendous apical excess surface area. As a consequence, neighboring WT cells are stretched out and respond by surface area dilatation. C) To test the relative impact of adhesion and contractility, the latter was balanced again by drug addition, revealing a temporal dependency: balanced contractility restores random mixing at early stages but differential adhesion is still able to promote cell sorting into clusters on long time scales.

It is well established that tight junction-depleted cells show increased contractility.(28, 31, 32, 36–38) However, so far the implications of increased contractility of dKD cells for the behavior of the monolayer was only studied with emphasis on impaired migration dynamics and signaling.(23, 24) Here, we showed that epithelial cells, which are stretched by their contractile neighbors respond primarily by apical area dilatation, instead of adaption of cortex mechanics. By comparing the apparent area compressibility modules of dKD and WT cells, we found a six-fold larger excess area for highly contractile dKD cells compared with dilated WT neighbors, equivalent to the change in geometric surface area, indicating the conservation of excess material instead of its recycling. This is consistently observed for co-cultures as well as dKD mono-cultures, which were described previously, where two populations emerged in a tug of war: a contractile population that stretches out the neighboring cell population to conserve the amount of occupied surface area.(23) However, in that work it remained unclear if recruiting excess surface area indeed dominates the stretch response. During development, a generation of two mechanical cell populations among the same cell type was identified as an emergent property upon collective cell interactions.(39) A recent study implicated asymmetric ROCK signaling in inducing these two populations to interact in confluent dKD mono-cultures.(24) This tug of war might intuitively favor segregation into clusters to decrease the number of WT cells that are subject to dilatation by adjacent dKD neighbors.

Our study confirms that segregation can be explained by the different interfacial tension model.(8) However, we found a temporal separation of the dominant source for demixing, where differences in contractility determine the cell sorting process on short time scales immediately after seeding, while differential adhesion contributes less but also permanently to the sorting process on longer time scales. On the one hand, adhesion-based sorting was shown before to emerge in cell cultures, e.g., upon different expression levels of cadherins.(6, 40–42) Similarly, sorting based on cadherin levels was demonstrated in follicle and retina cells of Drosophila oocytes.(43, 44) Signaling-controlled cadherin turnover has also been implicated in cell segregation.(45, 46) Note, in our study, adhesion differences were induced by tight junction disruption, which was also shown by previous work to decrease adhesion, in agreement with our data.(30, 47) Purely adhesion-based sorting was recently confirmed via simulations and experiments in direct relation to constant contractility.(18) On the other hand, differential contractility was found to aid sorting in an embryo and possibly dominate over adhesion.(48, 49) In co-cultures of zebrafish germ layer cells, differential contractility alone was found to be sufficient for sorting.(48) However, while sorting also took place on two time scales, a fast, early (< 0.5 h) and a slower, later time scale, the authors did not investigate the temporal evolution further. An interplay between adhesion and contractility was found in cancer cell line aggregates,(50) and confirmed in recent studies using vertex/Voronoi models.(9, 15, 16) In particular, interfacial tension was shown to be determined by the ratio of cell adhesion and contractility, governing the tissue-scale tension.(9) Accordingly, the increased contractility paired with the lower adhesion of dKD cells translates well into the high tension values measured by laser ablation.

However, a demixing mechanism of locally increased contractility at the boundary between two cell types, as reported in Drosophila wing discs, can be ruled out in our work.(10–12) While we measured tremendous differences in line tension between the different cell types, the WT-GFP/dKD interface did not exhibit the highest tension but rather values equal to or slightly below that of dKD/dKD junctions.

Another mechanism in contrast to our data was proposed by a recent study examining the demixing of E-cadherin-depleted and wildtype MDCK cells. The authors identified active cell forces as the governing factor of sorting.(13) While the demixing behavior in that study appears very similar to our data, the initial segregation was slower. Furthermore, they observed a pattern reversal at uneven mixing ratios which were absent in our co-cultures. This stability of the sorting pattern is indicative of interfacial energy minimization by minimizing the contact region between heterotypic cell types upon sorting, based on adhesion and/or contractility.(13) E-cadherin-depleted and wildtype keratinocytes were recently shown to sort mainly based on shape disparities and this was thoroughly explained in vertex simulations as well as observed earlier in zebrafish embryos.(14, 51) Although the shape differences in our cell lines seem to be small and result from the tug-of-war between the cell types, we cannot entirely exclude their contribution.(23)

We also addressed the possible crosstalk of contractility and adhesion.(52) For reference, actomyosin contractility has been shown to enhance adherens junction-based adhesion.(53–55) Measurement of cell-cell adhesion forces in the presence of Y27632 showed that albeit adhesion was slightly reduced due to loss of contractility, the difference between the forces measured for WT-GFP/WT and WT-GFP/dKD cell pairs, respectively, remained unchanged.

However, there would still be the disruption of the tight junctions in dKD cells. In addition, if the adhesion difference had been just as abolished as the differential contractility, we would not have observed the prevailing demixing at longer time scales. Actomyosin contractility has also been shown to modulate focal adhesions and thereby cell motility.(56–58) However, we observed no influence of motility on sorting, possibly due to the high cell density in our experiments (with confluence reached after only a few hours).

In addition, cell sorting could be influenced by proliferation creating local clusters. However, upon Y27632 treatment the WT-GFP/WT control increased its proliferation to the same level present in treated and untreated WT-GFP/dKD mixtures, yet, its *SI* remained much lower. At the same cell amount, the *SI* remained lower in the treated WT-GFP/WT control than in the untreated WT-GFP/dKD mixture. In addition, the large difference in proliferation between the treated and untreated WT-GFP/WT samples did not translate into an increase in segregation. The cell amount ratio of the cell types was also consistent among the WT-GFP/dKD mixture and its respective WT-GFP/WT counterpart, both treated and untreated, whereas their *SI* differed. Ultimately, using mitomycin C to suppress proliferation we found indeed that the early stage of cell sorting dominated by differences in contractility was unaltered.

Another caveat to note is that due to technical limitations, our cell adhesion measurements are on much shorter timescales than the observed mixing dynamics and the relevant cell-cell interactions in general.(59) Nevertheless, in agreement with other work, it is reasonable to assume that the loss of tight junction integrity reduces intercellular adhesion on all relevant timescales.(30, 47) While ZO proteins do not directly bind to the other cell, their loss destabilizes the contact zone and influences the transmembrane proteins.(30, 60)

In general, the presented model system has the advantage of great experimental accessibility compared with in vivo experiments but lacks some physiological conditions, e.g., properties of the substrate. While the cell-substrate adhesion might be different, it is well suited to capture the general physics of cell-cell interactions that also governs sorting in vivo. Along this line, previous work successfully compared consistently in vivo and in vitro tension-based cell sorting experiments.(48, 49) Ultimately, our data suggest that adhesion alone is sufficient but less efficient in driving cell sorting without differential contractility. This could yet be another example of how biology employs functional redundancy to ensure fundamental processes such as the sorting of different cell types.

## Materials and Methods

Full materials and methods are available in the SI section. MDCKII cells were used for all experiments, and knockdowns or fluorescent protein labeling were performed using Crispr/Cas as described in Beutel et al.(61) or Skamrahl et al.(23) For the figures, images were brightness-adjusted in Fiji to improve visibility.(62) Segmentation was performed using Cellpose 1.0.(63) Cell positions and areas were obtained using OpenCV as described in Skamrahl et al.(23, 64, 65) Neighbor analysis was performed with self-written Python scripts. Trackpy was used for tracking.(66, 67) Confocal microscopy (FluoView1200; Olympus, Tokyo, Japan) was performed after fixation and antibody- and phalloidin-based labeling. Analysis was performed using segmentation via Cellpose. AFM was carried out on a NanoWizard 4XP (Bruker Nano, JPK, Berlin, Germany) and calibrated by the thermal noise method.(68) Imaging was performed in contact mode after fixation. Indentation curves were analyzed as described recently(23) using the viscoelastic theory introduced by Cordes et al.(26) An 800 nm femtosecond pulsed laser was used for laser ablation and the opposing vertex knots of a junction were tracked. Cell-cell adhesion measurements were performed on a Cellhesion 200 AFM (JPK) at 0.5 μm s^-1^ (single cells) / 1.0 μm s^-1^ (cells grown on cantilever) and the peak adhesion force was evaluated. Cells were cultured on collagen-coated cantilever. Proliferation was inhibited by incubation for 1 h with 10 μg/mL mitomycin C which was removed by centrifugation before seeding. Calcium withdrawal was performed using minimum essential medium eagle, spinner modification (SMEM; Sigma-Aldrich life science, UK) and media were supplemented with chelex serum (FCS Gold neutralized chelex treated, PPA, Pasching, Germany) to better control the calcium concentration. Cluster analysis was performed on the WT-GFP fluorescence images using scikit image in Python.

## Acknowledgments

Funding from the DFG grants SPP1782 and DFG JA963/19-1 is gratefully acknowledged. We thank Burkhard Geil and Jonathan F.E. Bodenschatz for helpful discussions.

