## Supporting Information for "Cellular segregation in co-cultures driven by differential adhesion and contractility on distinct time scales"

Andreas Janshoff

**This PDF file includes:**

Supporting Materials and Methods

Figures S1 to S11

### Materials and Methods

#### *Cell culture handling*

Madin-Darby Canine Kidney cells (strain II, MDCKII; European Collection of Authenticated Cell Cultures, Salisbury, UK) were cultured at 37°C, 5% $CO_2$, humid conditions, and in minimum essential medium (Life Technologies, Paisley, UK) containing Earle's salts, 2 mM GlutaMAX (ThermoFisher Scientific, Waltham, Massachusetts, USA), 2.2 g $L^{-1}$ $NaHCO_3$, and 10% fetal bovine serum (BioWest, Nuaillé, France), here termed $M10F^-$. The cells were passaged two to three times per week before reaching confluence with phosphate buffered saline pH 7.4 (PBS; Biochrom, Berlin, Germany) containing trypsin/EDTA (0.25%/0.02% w/v; BioWest/Biochrom).

*Genetic generation of cell lines*

ZO1/2 knockdown was effected according to Beutel et al. using Crispr/Cas.(61) WT-GFP cells were created as described in Skamrahl et al.(23) Clones of MDCKII cells expressing GFP-myosin-2-A were generated by transfecting cells with pTRA-GFP-NMCH II-A plasmid (Addgene plasmid # 10844). Clones expressing the GFP tag in a stable manner were selected via neomycin resistance (G418). Upon selection, the cell pool was sorted using FACS to enrich cells with GFP at a moderate level. For N-terminal endogenous labeling of myosin with mNeon in dKD cells, the myosin-2-A exon was targeted using Crispr/Cas. MDCKII WT-ZO1-mNeonGreen cells were generated as described by Beutel et al. targeting the initial ZO1 exon with Crispr/Cas.(61)

*Cell seeding and demixing experiments*

Petri dishes (µ-dish, 35 mm, ibiTreat 1.5 polymer surface; ibidi, Martinsried, Germany) were used. Cells were trypsinized and mixed well before being seeded to ensure an initially random distribution.

Two sets of seeding conditions were used: In *seeding approach one*, cells were seeded at $6 \times 10^5$ cells in 1 mL $M10F^-$, rinsed after about 5 h with $M10F^-$, supplied with sufficient $M10F^-$ (2-3 mL), and then imaged over time or fixed/measured after 28 h or 48 h. These conditions were used for the experiments shown in Figure 2 and 3. Because in the first experimental approach the initial demixing dynamics were missed and the cells were still subconfluent for a long time (while the image quality was slightly better due to the rinsing step and the lower cell density), we changed the experiment: Cells were seeded at $1.2 \times 10^6$ cells in 1 mL $M10F^-$ and imaged subsequently. This *second seeding approach* was used for the main demixing experiments shown in Figure 1, 5 and S1-3. To reduce cellular contractility, the same second set of experiments was performed with Y27632 (Y27632; Sigma-Aldrich, Steinheim, Germany) added to the mixed cell solution, to reach the desired final concentration, immediately before seeding. Importantly, the cell parameters (*SI*, cell area ratio, relative cell amount as well as morphology) were almost identical for both seeding conditions.

For imaging, cells were placed into the incubation system of a fully automated inverted light microscope (BZ-X810; Keyence, Neu-Isenburg, Germany) equipped with a 10x phase contrast objective (Nikon CFI60 Series; Keyence). The temperature was calibrated to be 37°C at the cells using a local temperature probe (Testo 735; Testo, Lenzkirch, Germany), 5% partial $CO_2$ pressure was set, and sufficient humidity was provided with distilled water in the appliance of the incubation system as described before.(23) Phase contrast images were recorded at 1 frame per 7.5 min, 14 bit, 25% illumination power, exposure times of 1/25 s, and without binning, zoom, or gain, yielding a field of view of 1920 x 1440 pixels (1449.6 µm x 1087.2 µm). Corresponding fluorescence images for each frame were recorded at low light exposure to prevent phototoxicity. For this, we used 40% illumination power, 3.5 s exposure and 4x gain, without binning to allow for direct overlay of both channels. In between frames, all light exposure was turned off. Focus tracking was applied and three vertical slices were chosen in a range of 10 µm (5 µm pitch) to avoid drift effects. For the manuscript figures, images were brightness-adjusted in ImageJ (Fiji) to improve visibility,(62) particularly in relation to the phase contrast. The fluorescence brightness was typically increased (images were relatively dark due to the low-exposure settings).

*Demixing experiments with mitomycin C-inhibited proliferation*

Mitomycin C (MitoC; Sigma-Aldrich, Steinheim, Germany) was dissolved in water to reach 500 µg $mL^{-1}$ and stored in aliquots of 150 µL. Co-culture solution was prepared at 1:1-mixing ratio in PBS with 10 µg/mL MitoC in an incubator at 37°C, 5% $CO_2$ for 55 min. After thorough mixing, 0.25 % trypsin in PBS (w/v, BioWest) was added to isolate cells and cells were incubated for 5 min. Then, to remove the MitoC and prevent longterm cytotoxic effects, the cells were centrifuged at 1200 rpm for 3 min to aspirate the supernatant and resuspend the cell pellet in 1 mL fresh M10F-. Lastly, $1.2 \times 10^6$ cells in 1 mL M10F- were evenly distributed on µ-dishes (µ-dish, 35 mm low, ibiTreat No. 1.5 polymer surface; ibidi) and imaged immediately as described above. Controls were performed accordingly but without MitoC addition.

*Cell sorting experiments with varying calcium concentrations*

For the $Ca^{2+}$ withdrawal assay experiments, different solutions with variable $Ca^{2+}$-concentration were used: the regular $Ca^{2+}$ medium (minimum essential medium containing Earle's salts, 2 mM GlutaMAX, and 2.2 g $L^{-1}$ $NaHCO_3$, as described above for all other experiments), a $Ca^{2+}$-free-medium (minimum essential medium eagle spinner modification (SMEM; Sigma-Aldrich life science, UK) containing Earle's salts, 2 mM glutamine, and 2.2 g $L^{-1}$ $NaHCO_3$) and a mixture of these two media to realize specific $Ca^{2+}$-concentrations. Media were supplemented with chelex serum (FCS Gold neutralized chelex treated, PPA, Pasching, Germany) to better control the final calcium concentration.

For co-culture in the presence of calcium, cells were passaged and resuspended in 1 mL of regular $Ca^{2+}$ medium. Thereafter, $6x10^5$ cells/mL of each cell line, i.e., in total $1.2x10^6$ cells/mL were seeded evenly on µ-dishes (35 mm, ibiTreat 1.5 polymer surface, ibidi). In the experiments with $Ca^{2+}$-free and $Ca^{2+}$-depleted medium, cells were resuspended in $Ca^{2+}$-free medium, and the $Ca^{2+}$ concentration was adjusted by adding a defined fraction of the regular $Ca^{2+}$ medium before seeding.

*Automated cellular segmentation*

Cell segmentation was performed as described in Skamrahl et al.(23) using Cellpose 1.0(63) in conjunction with Python-based parallel processing. Raw phase contrast or fluorescence images were directly used as input. To optimize cell recognition, the following parameters were used: for the phase contrast channel, the flow and cell probability thresholds were set to 1 and -6, respectively. For the fluorescence channel, the flow and cell probability thresholds were set to 0.9 and -5, respectively.

*Further automated analysis of cell parameters*

To calculate cell parameters such as the x- and y-position, area, and aspect ratio, OpenCV was used as described in Skamrahl et al.(23, 64, 65) Note that OpenCV might omit a small amount of cells present in the Cellpose data, for example due to a failed aspect ratio calculation (e.g., due to a falsely recognized particle which is only a few pixels in size). Therefore, to ensure correct cell indexing between Cellpose and OpenCV in all functions, we used the contours (and masks) from OpenCV for all following operations. For cell tracking, Trackpy(66) was used to link the cell positions from OpenCV. The link function was used with a memory of 3 frames and 6.04 µm

(8 pixels) as the maximal displacement. Trajectories shorter than 5 frames were discarded. Drift correction was not necessary.

*Automated cell type recognition, neighbor analysis, and segregation index calculation*

Using the masks generated by Cellpose, three main steps were performed in Python. To identify the cell type (WT-GFP or unlabeled WT in WT-GFP/WT, WT-GFP or unlabeled dKD in WT-GFP/dKD mixes), the mask returned by Cellpose for the fluorescence and phase contrast channel were compared. In the masks, all pixel values belonging to one cell body correspond to the respective cell index with the background being zero. We iterated over the masks in the phase contrast channel and evaluated the modal value of the respective cell pixels in the fluorescence channel. If the modal value of the respective cell pixels corresponded to zero in the fluorescence channel (i.e., background without a cell), the cell was assigned as unlabeled. If the modal value was greater than zero, it was assigned as WT-GFP. The cell types were assigned in the phase contrast masks to finally yield a complete set of all cells for each image. Next, nearest neighbor analysis was performed to find direct neighbors. For each cell, an OpenCV line scan was performed between the center of this cell and each surrounding cell. All zeros, the own cell index, and the index of the respective neighbor were removed from the line. Only if no values remained in the line, i.e., no other cell index was crossed, the two cells were assigned as neighbors. This was restricted to a window of 50 pixels x 50 pixels (37.75 µm x 37.75 µm) around each cell to prevent extremely high computation times and also false neighbor assignment of cells that are separated by empty space in the early, not fully confluent state. Neighbor assignments were noted as ones in a so-called adjacency matrix, in which the column (or row) number corresponds to the cell index, the rest of this symmetric matrix was filled with zeros. Lastly, the number of neighbors of each cell in the image was the sum of all ones in the respective column (or row) in the matrix. We distinguished the number of homotypic neighbors by only summing over the positions in the matrix given by the cell assignments. Per definition, the segregation index *SI* was the ratio of the homotypic and all neighbors for each cell. Finally, to generate the plots, the *SI* was averaged over all cells or separately over all cells of each cell type.

*Cluster analysis of unlabeled cells (WT or dKD, respectively)*

The properties of the unlabeled cell clusters were quantified directly from the WT-GFP fluorescence time lapse images using Python's skimage modules.(69) First, images were normalized and brightness adjusted using exposure.equalize_adapthist with a clip limit of 0.04

and 256 bins. The images were thresholded below 0.25 to remove the bright WT-GFP cells. Small "objects" below 200 pixels and "holes" below 10 pixels, such as the dark nuclei, were removed. Finally, measure.label and .regionprops were used to extract all cluster properties.

*Cell labeling and confocal fluorescence microscopy*

For the depicted confocal images, 6 $10^5$ cells in 1 mL were seeded (first seeding approach, *vide supra*) and fixed after 28 h or 48 h. Before labeling, samples were incubated for 20 min with a paraformaldehyde/glutaraldehyde solution (4% (w/v) / 0.1% (w/v) in PBS; Science Services, Munich, Germany / Sigma-Aldrich). Cells were incubated for 5 min in Triton X-100 (0.1% (v/v) in PBS) to permeabilize the plasma membrane. After rinsing three times with PBS, samples were incubated in blocking/dilution buffer (PBS containing 2% (w/v) bovine serum albumin and 0.1% (v/v) Tween20) for 30 min to block unspecific binding sites.

Samples were incubated in primary antibody diluted in blocking/dilution buffer for 1 h using the following reagents. Phospho-myosin: 2 µg $mL^{-1}$ (1:200) light chain 2 (Ser 19) rabbit IgG1 (Cell Signaling Technology, Danvers, Massachusetts, USA), ZO1: 5 µg $mL^{-1}$ (1:100) mouse ZO1-1A12 IgG1 AlexaFluor 488 (Invitrogen, ThermoFisher Scientific, Waltham, Massachusetts, USA). After incubation with the primary antibody, samples were briefly rinsed with PBS. Next, they were washed with PBS, with 0.1% (v/v) Triton X-100 in PBS, and again with PBS, each for 5 min on a shaking plate (75 rpm).

The secondary antibody (AlexaFluor 546-conjugated goat anti-rabbit IgG; Life Technologies, Carlsbad, USA) was diluted with blocking/dilution buffer to a concentration of 5 µg $mL^{-1}$. Actin labeling was performed using 165 nM AlexaFluor 647-phalloidin (Invitrogen), incubated and diluted together with the secondary antibody. The incubation time was 1 h. Following the secondary antibody, samples were washed three times with PBS for 5 min each on a shaker (75 rpm).

Nucleus staining was performed by a 15 min-incubation with DAPI (Sigma-Aldrich), diluted to 50 ng $mL^{-1}$. Before imaging, the cells were rinsed with PBS three times and then kept in PBS. Labeling and microscopy were performed at room temperature. For live-cell nuclei staining of WT-GFP cells cultured on an AFM cantilever (Figure S11), Hoechst 33342 (Molecular Probes, Thermo Fisher) was added at 1 µg/mL in $M10F^+$ and incubated for 30 min in the incubator. Afterwards it was rinsed with $M10F^+$ and $M10F^+$ was used during imaging.

For fluorescence imaging experiments a confocal laser scanning microscope (FluoView1200; Olympus, Tokyo, Japan) was used with a 60X objective (oil immersion, $NA = 1.25$). Image processing (3D representations, z-projections, color choice, and overlay) and brightness adjustment were performed in Fiji.(62) For the figures, brightness usually had to be slightly increased due to low-bleaching acquisition settings.

*Fluorescence image intensity quantification*

Both, the P-myosin-2 and the actin signal were quantified using Cellpose-based segmentation. Thresholds were manually adjusted and the Cellpose GUI was used to improve the segmentations. The masks from Cellpose were then used to generate cell centers and junctions using OpenCV after subtracting the outlines. The ZO1 centers (WT cells only) were used to distinguish between WT and dKD cells applying the same cell type recognition strategy as described above with a search window of 100 pixels. After the cell types were detected, 5-pixel thick outlines of the junctions were subtracted from the center masks. In addition, the center masks had to be set to zero at the positions of the original (1-pixel thin) outlines. After these steps, the center masks are actually reduced cell centers without the junction signals. Finally, for each cell, the intensities of the original images at the 5-pixel thick junctions were averaged and divided by the averaged intensities from the centers to determine the intensity ratios Figure 2 A2). Absolute intensities were calculated from the original OpenCV center masks and averaged for each cell. Note, at the apical plane the masks were obtained from maximum intensity projections (Fiji) of the actin signal. In this instance, projections were used to capture all cells by equalizing the height difference between dKD and WT (otherwise WT cells would be barely visible).

*AFM imaging*

Two days after seeding at 6 $10^5$ cells in 1 mL (first seeding approach), cells were rinsed once with PBS containing 0.1 g $L^{-1}$ $Mg^{2+}$ and 0.133 g $L^{-1}$ $Ca^{2+}$ ($PBS^{++}$; Sigma-Aldrich) and incubated with glutaraldehyde solution (2.5% (v/v) in $PBS^{++}$) for 20 min. $PBS^{++}$ was used instead of PBS without magnesium and calcium ions, because the dKD cell layers were more susceptible to dissolution of ion-dependent adhesion sites due to the impaired diffusion barrier function. Prior to AFM imaging, samples were rinsed three times to remove residual glutaraldehyde. A NanoWizard 4XP AFM (Bruker Nano, JPK, Berlin, Germany) was used for imaging. The AFM was mounted on an inverted light microscope (IX 83; Olympus) to allow visual inspection via phase contrast and cell recognition via fluorescence. Imaging was carried out in contact mode with MSCT C cantilevers

(Bruker AFM Probes, Camarillo, USA) in PBS with a line scan rate of 0.18 Hz translating to a velocity of 40 µm $s^{-1}$, a force of 0.14 nN, and a pixel size of 50 nm. The IGain was set to 20 Hz and the PGain was 0.002. The AFM was calibrated as described below, in a small area at the edge of the dish where a few cells were scratched off. Height and error images were obtained from the manufacturer's SPM Data Processing software (JPK, Berlin, Germany) upon standard linear plane correction (to correct for a slight tilt of the Petri dish surface) without further processing.

*AFM indentation and force relaxation* measurements

Force indentation-relaxation experiments were performed using a NanoWizard 4XP AFM (Bruker Nano) with a 37°C-heated stage (JPK) mounted on an inverted microscope (IX 83; Olympus) using silicon nitride cantilevers with a nominal spring constant of 0.01 N $m^{-1}$ (MLCT C; Bruker AFM Probes). Prior to experimentation, cantilevers were rinsed with isopropanol and PBS and functionalized by incubation for 1 h with fluorescein isothiocyanate-conjugated Concanavalin A (2.5 mg $mL^{-1}$ in PBS; Sigma-Aldrich).

The sensitivity of the AFM was determined by recording force curves on bare substrate and the spring constant of each cantilever was determined by the thermal noise method.(68) To allow the co-culture system to fully establish its differential collective mechanics, we allowed 2 days before measuring (with the first seeding approach). Cells were rinsed three times with $M10F^-$ containing 0.2 mg $mL^{-1}$ Penicillin (Biochrom), 0.2 mg $mL^{-1}$ Streptomycin (Biochrom), and 15 mM HEPES ($M10F^+$; BioWest).

Before the experiments, 2.5 mL $M10F^+$ was supplied to the cells and the temperature was set to 37°C. The cells were indented at a constant speed of 2 µm $s^{-1}$ to a maximum force of 1 nN. After a dwell time of 1 s at constant height the indenter was retracted at the same speed. Maps were recorded at a lateral scan resolution of 2 µm per pixel with one indentation-relaxation cycle each. Furthermore, five consecutive force curves at the center of individual cells in the monolayer were measured with the same parameters. For the data in Figure 3B

the individual curves at the cell center were pooled with curves at the cell center from the maps.

*Force curve fitting and viscoelastic model (Evans model)*

Indentation-relaxation curves of the cell center were analyzed as described recently(23) using the viscoelastic theory introduced previously by Cordes et al.(26) Briefly, the cell surface is described as a spherical cap and the cell is considered as a liquid-filled object surrounded by a thin isotropic viscoelastic shell, which is deformed at constant volume. We assume that the cell is surrounded by an isotropic elastic shell that produces a restoring force in response to indentation with a conical indenter originating only from two sources, linear elasticity due to area dilatation at large strains and pre-stress (constant homogenous tension) stored in the membrane and actin cortex. Pre-stress of the cortex/membrane composite is mainly generated by active elements (actomyosin), adhesion to the surface, and adhesion to the plasma membrane. Bending and shearing of the cell cortex are neglected in the following treatment since the cortex is assumed to be thin (100-200 nm) compared to the radius of the cell. As usual in linear elasticity, the lateral tension of the plasma membrane in the absence of shear can be written as a 2D Hookean law to first order:

$$T = T_0 + K_\mathrm{A}\frac{A_\mathrm{n}-A_0}{A_0}, \qquad (1)$$

in which $T_0$ comprises cortical tension $T_\mathrm{c}$ of the actomyosin cortex, and $T_t$ the membrane tension including elements of adhesion to the cytoskeleton and in–plane tension of the membrane.

$$T_0 = T_\mathrm{c} + T_\mathrm{t} \qquad (2)$$

$K_\mathrm{A}$ is the area compressibility modulus of the plasma membrane, and $A_\mathrm{n}-A_0$ is the difference between the actual area $A_\mathrm{n}$ after compression and the initial area before compression $A_0$. Static equilibrium is captured by the Young-Laplace equation, which describes the pressure difference $\Delta P$ across the fluid interface as a function of surface tension $T$ and mean curvature $H$:

$$\Delta P = 2TH \qquad (3)$$

Assuming a constant curvature of the free membrane justified by a constant hydrostatic pressure difference and a constant isotropic tension, we arrive at

$$2H = \frac{\mathrm{d}u}{\mathrm{d}r} + \frac{u}{r} \qquad (4)$$

where $u = \sin\gamma$ serves as a shape function of the deformed object with $\gamma$, the angle between the surface normal to the vertical axis. Equation (4) is solved for the following boundary conditions:

$$\gamma = -\Theta\ (r = r_1)$$
$$\gamma = \phi\ (r = R_1),$$

with the radius $R_1$ at the base of the spherical cap and the contact angle $\phi$ upon deformation as defined before. $r_1$ is the contact radius with the conical indenter, $\theta = \frac{\pi}{2} - \vartheta$ with $\vartheta$, the cone half angle. Integration of equation (4) gives:

$$u = C_1 r + \frac{C_2}{r},$$

with two integration constants:

$$C_1 = \frac{R_1 \sin(\phi) + r_1 \sin(\Theta)}{R_1^2 - r_1^2}.$$
$$C_2 = -C_1 r_1^2 - r_1 \sin(\Theta)$$

Force balance at the base gives the response force *F* of the cell to deformation:

$$F = 2\pi \left( R_1{}^2 \left( \frac{R_1 \sin\phi + r_1 \sin\theta}{R_1{}^2 - r_1{}^2} \right) - R_1 \sin\phi \right) T(t). \quad (5)$$

The task is now to determine $\phi$ and $r_1$, which are sufficient to describe the shape of the indented cell. Besides force balance, the central assumption for this is the conservation of volume during the indentation process. The volume at any time can be computed from the shape as a function of indentation depth:

$$V = \int_{r_1}^{R_1} \pi r^2 \frac{\left(C_1 r + \frac{C_2}{r}\right)}{\sqrt{1-\left(C_1 r + \frac{C_2}{r}\right)^2}} \mathrm{d}r - \frac{\pi r_1^3}{3\tan(\theta)} \quad (6)$$

and be set equal to the volume of the cell prior to indentation ($V = (3R_0 - h)\frac{\pi h^2}{3}$), with *h* the initial height of the spherical cap before deformation). Since area dilatation of the cortex is the main source of the measured restoring force, we need to compute the actual area $A_n$ from the shape of the indented cap:

$$A_\mathrm{n} = \int_{r_1}^{R_1} \frac{2\pi r}{\sqrt{1-\left(C_1 r + \frac{C_2}{r}\right)^2}} \mathrm{d}r + \frac{\pi r_1^2}{\sin(\theta)} \quad (7)$$

The actual indentation depth can also be computed from the shape at *r* = 0:

$$\delta = h - \int_0^{r_1} \frac{u}{\sqrt{1-u^2}} \mathrm{d}r + r_1 \tan(\theta). \quad (8)$$

The procedure allows to obtain force-indentation curves $f(\delta)$ by computing the shape of the cell parameterized by $r_1$ and $\phi$. Now, a set of nonlinear equations for the shape of the deformed cell is solved to fulfill force balance and the boundary condition of constant volume. The resulting shapes are minimal surfaces to minimize the stretching energy. Viscoelasticity of the shell is included in the tension term *T*(*t*) of equation (5) through a time *t* dependent area compressibility modulus $K_\mathrm{A} = K_\mathrm{A}{}^0 (t/t_0)^{-\beta}$ with the scaling parameter $K_\mathrm{A}{}^0$ and $t_0$ = 1 s (set arbitrarily). Application of the elastic-viscoelastic-correspondence principle leads to the following hereditary integral for the overall tension:

$$T(t) = T_0 + \int_0^t K_\mathrm{A}^0 \left(\frac{t-\tau}{t_0}\right)^{-\beta} \frac{\partial \alpha(\tau)}{\partial \tau} \mathrm{d}\tau$$

with the areal strain $\alpha = (\Delta A/A_0)$. The shape of the deformed cap only needs to be computed once for one set of parameters ($R_1$, $\phi$, and $\theta$) and can be approximated with a polynomial to obtain analytically tractable heredity integrals. The same can be done to the areal strain. Polynomials to the order of 4 are usually sufficient for good accuracy. A piecewise function is fitted to the indentation part ($0 < t < t_m$) and relaxation part ($t > t_m$), respectively.

The average geometry prior to deformation was derived via AFM imaging, confocal microscopy, and AFM-combined phase contrast and fluorescence: for WT cells a radius of 12 µm and an initial cap angle of 15° were used, while in case of dKDs $R_1$ = 5 µm and an angle of 25° was used. Cells were chosen close to the WT-GFP/dKD interface to most closely compare the mechanical differential. Viscoelasticity of the shell is included in the tension term $T(t)$ of equation (5) through a time $t$ dependent area compressibility modulus $K_A = K_A{}^0(t/t_0)^{-\beta}$ with the scaling parameter $K_A{}^0$ and $t_0$ = 1 s (set arbitrarily). The difference in excess area $A_{ex}$ between cell types was calculated via the correction factor $K_A{}^0\ \frac{A_0+A_{ex}}{A_0}$ with the surface area $A_0$ of the cell cap as first described in a study comparing isolated membranes and epithelial cells.(27)

Self-written Python scripts in conjunction with the JPK SPM Data Processing software were used for the analysis. A linear fit prior to the contact point was applied to correct the baseline. JPK SPM Data Processing was used to determine the contact point. Poorly fitted curves, e.g., yielding non-physical (negative) parameters, were discarded.

*Traction force microscopy*

Glass Petri dishes (µ-dish, 35 mm, high, glass bottom, ibidi) were first treated with 150 µL 3-aminopropyltriethoxysilane (APTMS, abcr GmbH). After 5 min, the surface was washed three-times with distilled water and dried with nitrogen. Then, the glass surface was covered with 300 µL glutardialdehyde solution (2.5 % in PBS) for 30 min. Simultaneously, cleaned circular cover slips with a diameter of 13 mm (borosilicate glass; VWR, Darmstadt, Germany) were coated with dimethyldichlorosilane for 5 min. The coverslips were then washed with ethanol (70%) and distilled water and afterwards dried with a stream of nitrogen. 4 µL of orange fluorescent beads (diameter: 200 nm) at a concentration of 0.2% solids (FluoSpheres, carboxylate-modified microspheres, 505/515 nm; Thermo Fisher Scientific Inc.) were added to 246 µL of a previously mixed polyacrylamide solution (0.6 g acrylamide (AA), 0.006 g N,N'-methylenebisacrylamide (Bis-AA) in 8 mL distilled water). 2.5 µL ammonium peroxydisulfate (APS, 10 mg in 100 µL Milli-Q; Carl Roth GmbH, Karlsruhe, Germany) and 0.5 µL tetramethylethylenediamine (TEMED, Carl Roth GmbH) were supplemented to the solution and mixed. 5.2 µL of the solution was added to the prepared Petri dish, covered with a coverslip, and allowed to polymerize for 45 min. The coverslips were subsequently removed. The resulted gels were covered twice with 200 µL SULFO-Sanpah (1 mg in 2 mL water; ThermoFischer Scientific Inc.) and irradiated for 8 min with UV-Light (365 nm). Between each step, the gels were washed 2x with distilled water. Afterwards the gels were coated with collagen (1 µL of collagen I (bovine), 5 mg/mL, Gibco/Life Technologies) in 199 µL of 0.02 M acetic acid) for one hour at room temperature. The gels were stored for further use in distilled water at 4°C.

MDCKII WT and dKD cells were seeded on the gels for traction force microscopy experiments. On each gel, 80000 cells were seeded in 200 µL M10F$^-$. After two hours of incubation, 2 mL of M10F$^-$ was added to each gel. TFM experiments were performed the next day (approximately 24 h after seeding).

Images were taken with a confocal laser scanning microscope (Fluoview 1200, Olympus) equipped with a 60x oil immersion objective (UPLFLN 60XOIPH, Olympus). After the first image was obtained, trypsin (2.5 %) with 0.05 M EDTA (Invitrogen) was added and after 10 to 20 minutes, a second image was taken. The ImageJ plugin "Template Matching and Slice Alignment" was used to remove the drift of the sample during the measurement. A particle image velocimetry MATLAB application, PIVlab, was used to calculate the displacement field of the beads.(70, 71) The displacement was determined with four interrogation windows (256 px, 128 px, 64 px, 32 px), and forces were calculated subsequently from displacement fields.(72) A mesh size of 16, a Poisson ratio of 0.48, and the built-in manual noise selection were used.

*Laser ablation*

Laser ablation was performed on a Zeiss LSM 780 NLO system driven by Zen Black software version 11.00. The image pixel size was 0.268 µm x 0.268 µm. The objective used was a Zeiss C- Apochromat 40X/1.2 W. Ablation was performed with an 800 nm Titanium/Saphire femtosecond pulsed laser Chameleon from Coherent (Santa Clara, USA) with a power of 3.2 W at the laser head, 60% laser output set in Zen Black, reflected by MBS 760+, with pixel dwell time for photomanipulation of 7.2 µs, single iteration, ablation area was line scan, 10 pixels. For measuring the recoil velocity, the lateral membrane of MDCKII WT and MDCK ZO1/2 dKD cells was highlighted by ZO1-mNeonGreen and myosin-2-A-mNeon, respectively, with the following settings: mNeonGreen was excited with 488 nm (Argon Laser) with MBS 488/561/633, emission filter used was 490-570 nm, pixel dwell time 2.83 µs, approximately 7.7 fps with GaAsP detector.

Allowing the cell layer to fully establish its mechanics, laser ablation was performed after 2 days of growth (first seeding approach).

*Cell-cell adhesion measurements*

All cell-cell adhesion measurements were carried out using a Cellhesion 200 AFM (JPK). The AFM was mounted on an optical IX 83 microscope (Olympus) to allow the identification of GFP-tagged and unlabeled cells. Single cell experiments from Figure 4 were performed immediately after seeding in M10F$^+$. Prior, tip-less cantilevers (MLCT-O10 B; Bruker AFM Probes) were rinsed several times with distilled water and isopropanol and then treated with poly-D-lysine hydrobromide (0.1 mg mL$^{-1}$ in water, Sigma-Aldrich) for about two days. A cell was picked and centered as well as possible above a cell on the substrate. Indentation was performed to a force of 2 nN, and a dwell of 5 s at constant height was chosen. The approach and retract velocity were set to 0.5 µm s$^{-1}$. The experiments were restricted to 2 h per day to avoid proliferation and advancing adhesion, which could interfere with the measurements. The adhesion peak force was determined using the JPK Data Processing software after linear baseline correction and contact point detection.

For the adhesion measurements in Figure S11, 5000 cells in 70 µL M10F$^-$ were grown overnight in ibidi inserts (ibidi) on µ-dished (35 mm, ibiTreat 1.5 polymer surface, low; ibidi). Cantilevers with a spherical tip (15 µm diameter CP-PNPL-SiO-E; NanoAndMore, Wetzlar, Germany) were used and rinsed several times with distilled water and isopropanol (p.a.). To culture cells on the cantilever, the cantilever was first coated for 1.5 h with 50 µL of 50 µg mL$^{-1}$ collagen I (bovine, in 0.02 M acetic acid; Gibco/Life Technologies) dispensed directly on the back of the cantilever. Cells were seeded at the highest possible concentrations (3.5 – 4.5 $10^6$ cells mL$^{-1}$), dispensing 75 µL at the cantilever tip. After roughly 2 h, another 50 µL was added. Then, 2 mL of M10F$^+$ was added and cells were grown overnight. After removing the insert, cells on the dishes were washed once with M10F$^+$ containing HEPES and the dishes were then filled with 2.5 mL M10F$^+$ and mounted on the Cellhesion 200 / optical microscope (IX 81; Olympus) setup. For the experiments with Y27632, the drug was supplemented to all media, including the medium used for overnight growth. The cantilever was mounted and checked for cells at the tip using the WT-GFP fluorescence signal. Approach was performed at 5 µm s$^{-1}$, the delay in contact was at constant height for 5 s, and the retract was at 1 µm s$^{-1}$. For the experiments estimated calibration parameters were used initially. The AFM was then calibrated after experimentation and removal of cells with 2.5 % trypsin. Cell removal was checked optically before performing thermal calibration and indentation on bare substrate. The evaluated forces were adjusted accordingly, yielding indentation setpoints of 2.1 – 2.8 nN.

*Statistical analyses and reproducibility*

The cell behavior was very reproducible between different samples as well as among seeding conditions. The significance of the AFM indentation data in Figure 3 was tested using the Mann-Whitney U test. The cell velocity and persistence were tested with the student's t-test. The fluorescence intensities in Figure 2A2, the laser ablation data in Figure 3 and the cell-cell adhesion forces in Figure 4 were tested using Welch's t-test. All statistical analyses were performed in Python.

A *p-value* of $< 0.05$ was considered significant and denoted by one asterisk (*). $p < 0.01$ and $p < 0.001$ were indicated by two (**) and three (***) asterisks, respectively.

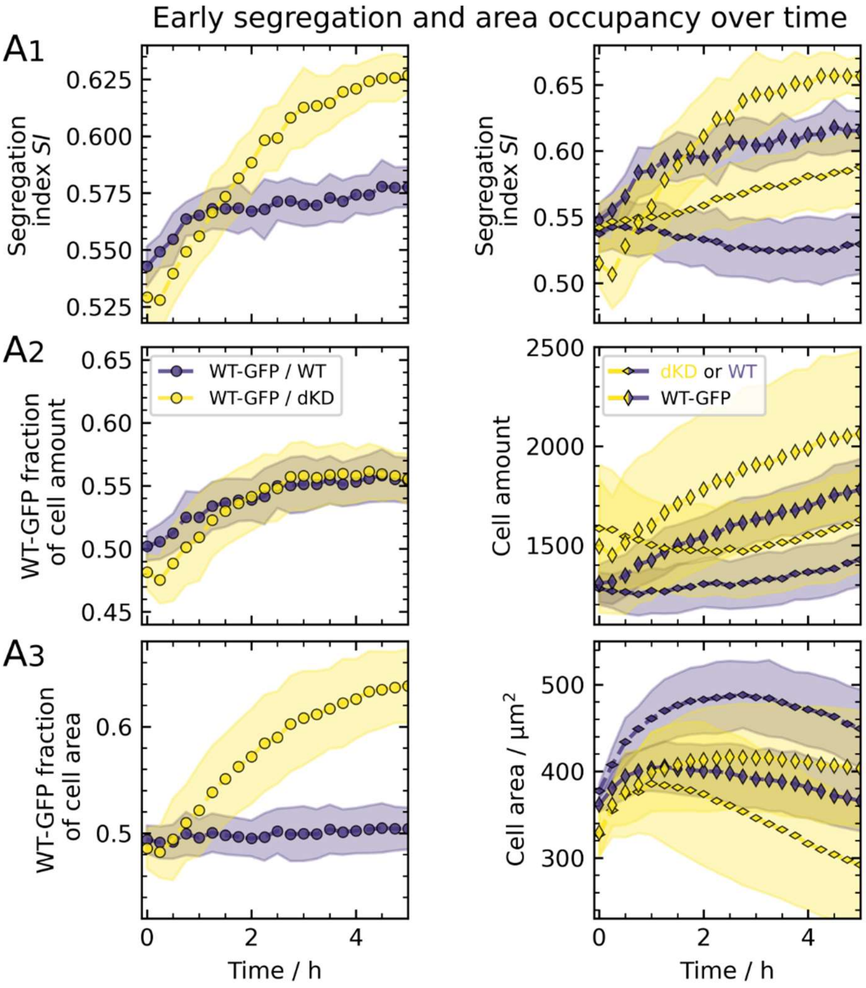


**Figure S1.** Early demixing behavior of dKD and WT cell co-cultures at an initial mixing ratio of 50:50, zoomed-in from Figure 1. All panels (A1-3) are set up as in Figure 1 and show the same data only with a limited time scale.

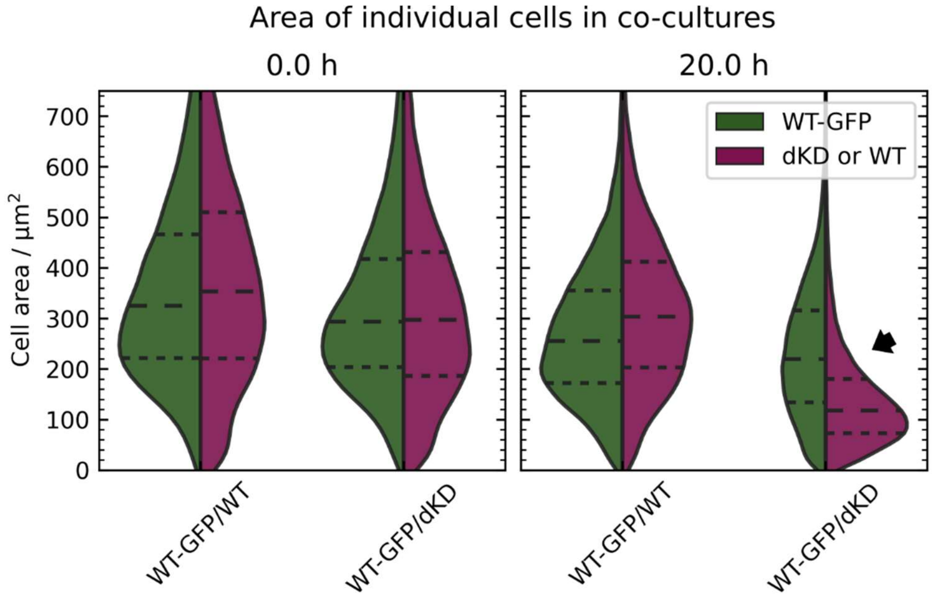


**Figure S2.** Area of individual cells in the co-cultures from Figure 1. Violin plots are shown, depicting the kernel density estimation with horizontal, dashed lines showing the quartiles and median. Violins are scaled to have the same area. The arrow indicates the unique area offset in WT-GFP/dKD co-cultures due to differential contractility.

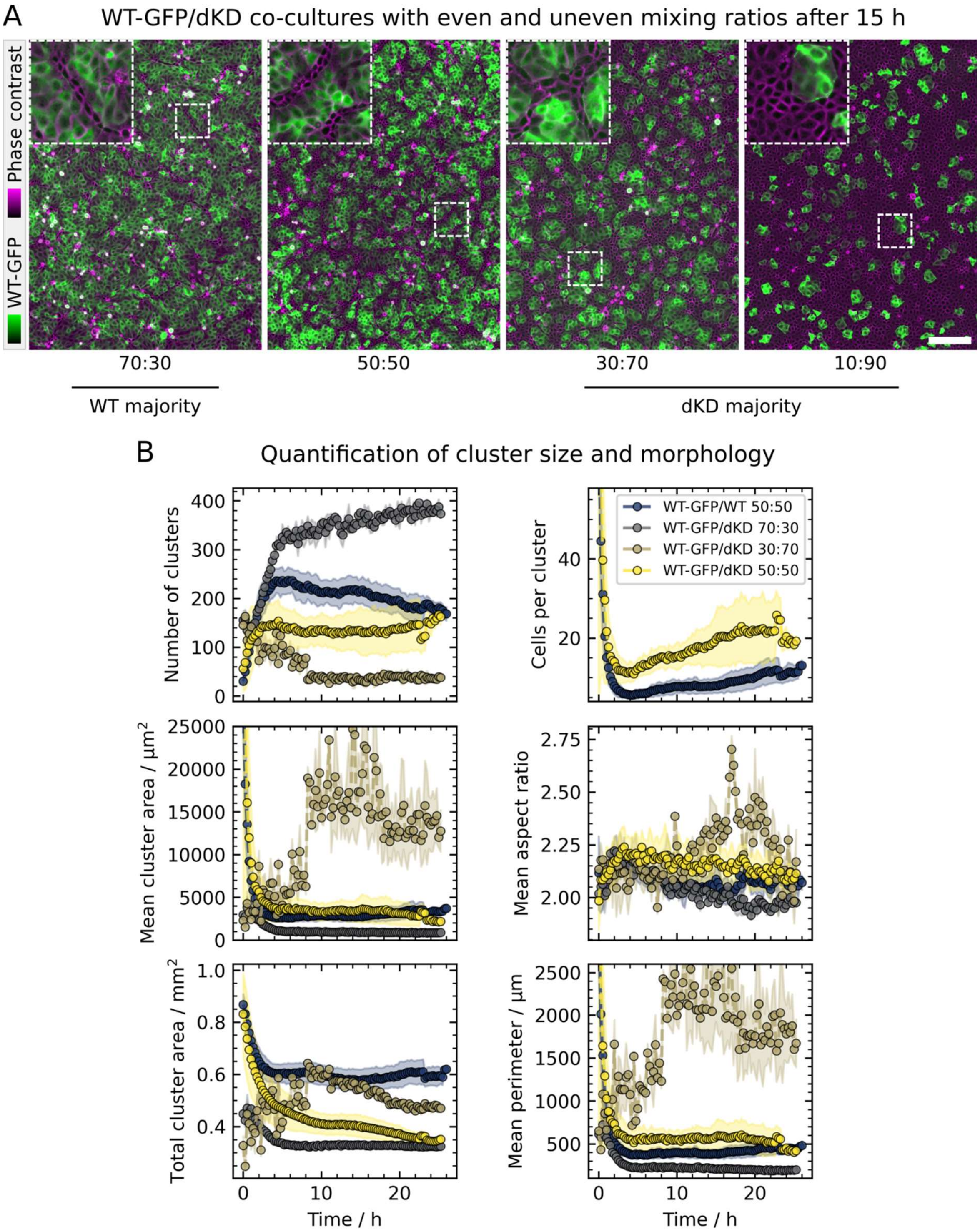


**Figure S3.** Demixing WT-GFP/dKD co-culture experiments performed as in Figure 1, seeded at even and uneven mixing ratios. A) Phase contrast (pseudo-colored in magenta) and WT-GFP fluorescence after 15 h of incubation. Scale bar: 200 µm. B) Quantification of cluster size and morphology of the respective unlabeled cells (WT or dKD) in the co-cultures over time.

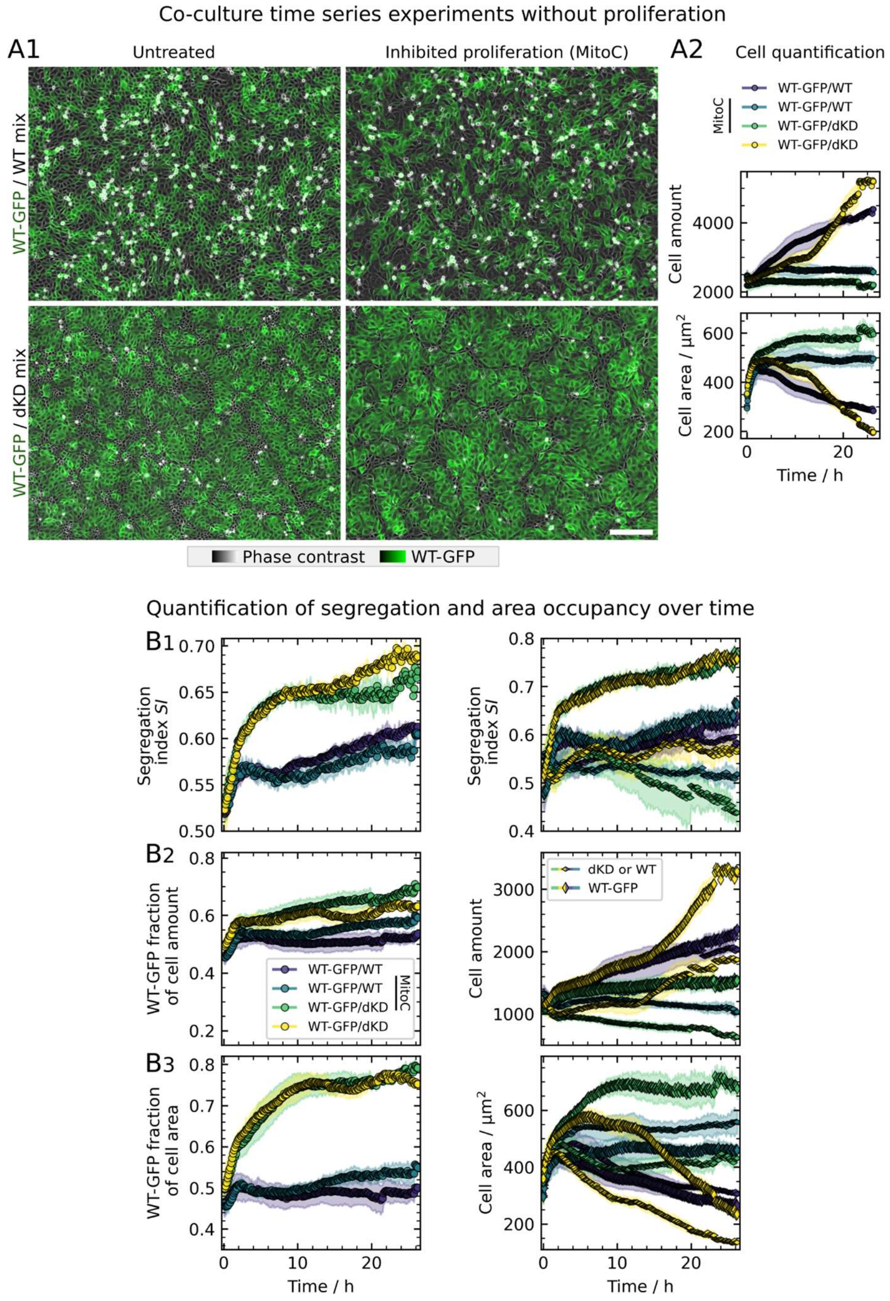


**Figure S4.** Proliferation does not play a major role in the segregation of WT-GFP/dKD co-cultures. A1) Example phase contrast images with superimposed WT-GFP fluorescence of co-cultures with and without proliferation (inhibited by mitomycin C (MitoC)) after 20 hours of growth. Scale bar: 200 μm. A2) Results of cell segmentation confirming successful inhibition of proliferation by MitoC. B1-3) Quantification parameters set up as in the B panel of the main Figures 1 and 5. B1 shows that proliferation does not impair cell sorting in the early stage of the process driven by differential contractility of dKD and WT cells, eventually reaching the same SI as in the unperturbed control sample.

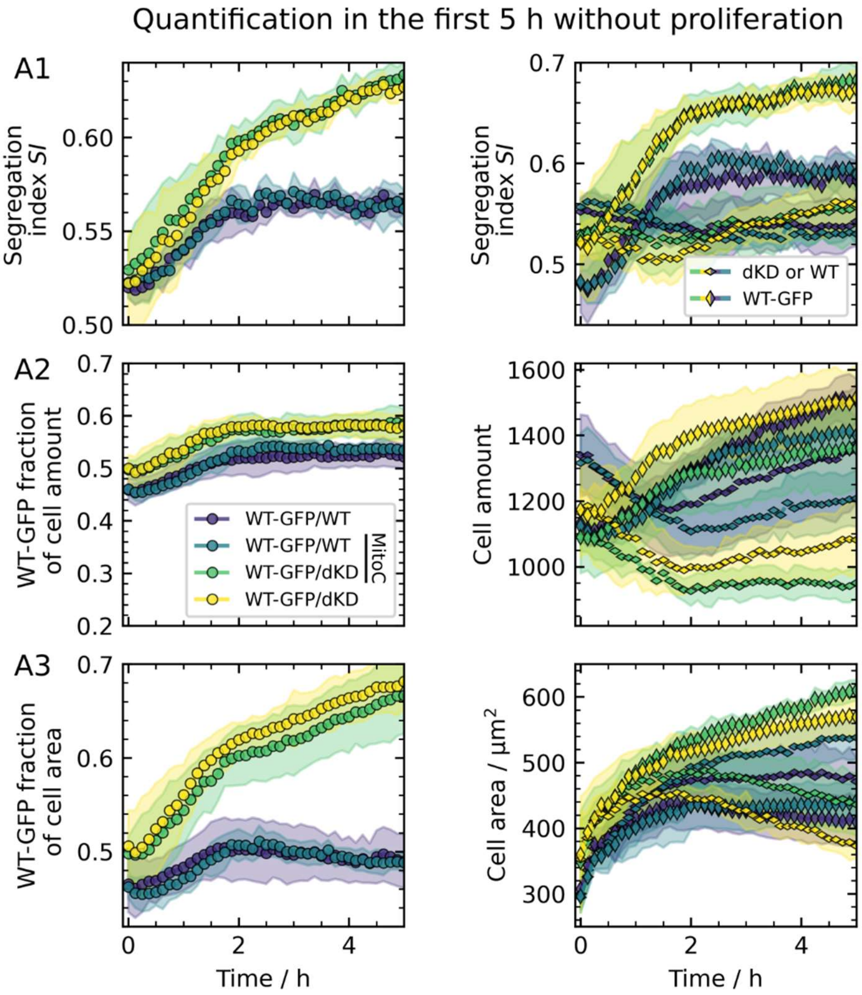


**Figure S5.** Zoom-in of the first 5the early demixing behavior of co-cultures at an initial mixing ratio of 50:50 with and without proliferation (inhibited by mitomycin C (MitoC)).

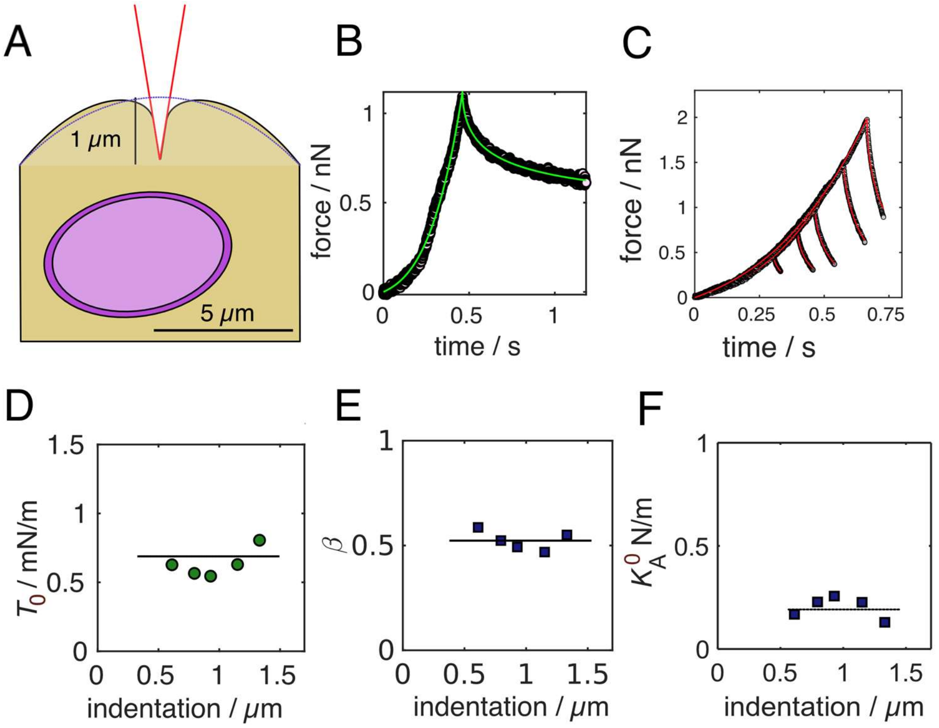


**Figure S6.** Impact of indentation depth on the mechanical response of confluent MDCKII cells. A) Scheme of the typical geometry of an MDCKII cell (WT) within a confluent monolayer. The approximate sizes of the cell and nucleus are inferred from CLSM images. B) Indentation-relaxation curve of a dKD cell indented with conical indenter and fitted with equation (5) (green line). C) The identical MDCKII cells (confluent WT cells) were indented with different yield forces leading to different indentation depths (velocity: 2 μm/s). Fits (red lines) of the viscoelastic Evans model provide almost identical parameters (pre-stress (D), fluidity (E), and area compressibility modules (F)) independent of the indentation depth. However, larger depths beyond 2 μm were avoided to exclude non-linear strain-stiffening effects as well as the impact from the substantially stiffer nucleus.(73, 74)

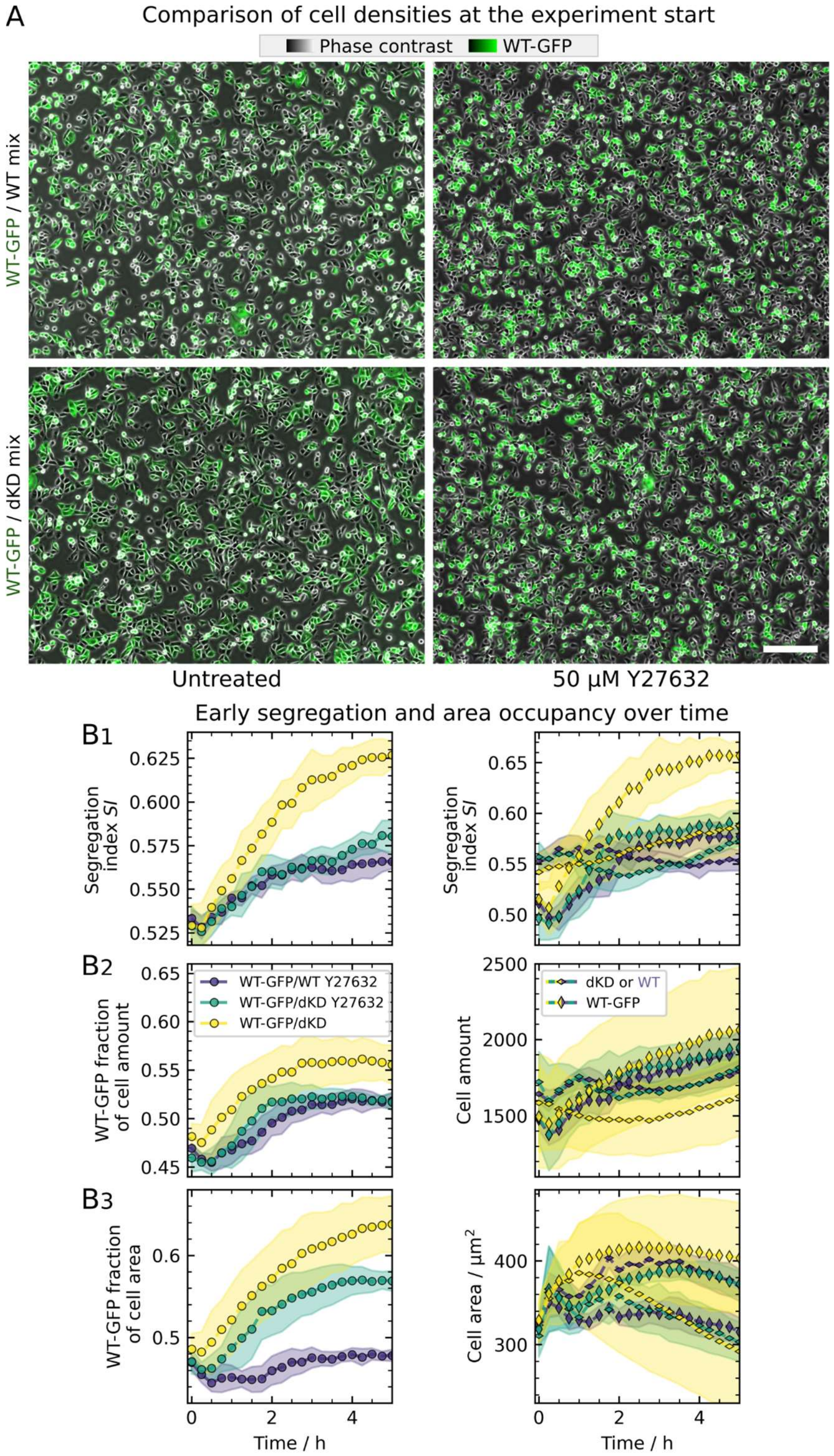


**Figure S7.** A) Comparison of different initial seeding densities at the experiment start (*t* = 0h) from Figure 1 and 5, respectively. B) Early demixing behavior of Y27632-treated dKD and WT cell co-cultures at an initial mixing ratio of 50:50, the zoomed-in data are taken from Figure 5. All panels (B1-3) are set up as in Figure 5B and show the same data.

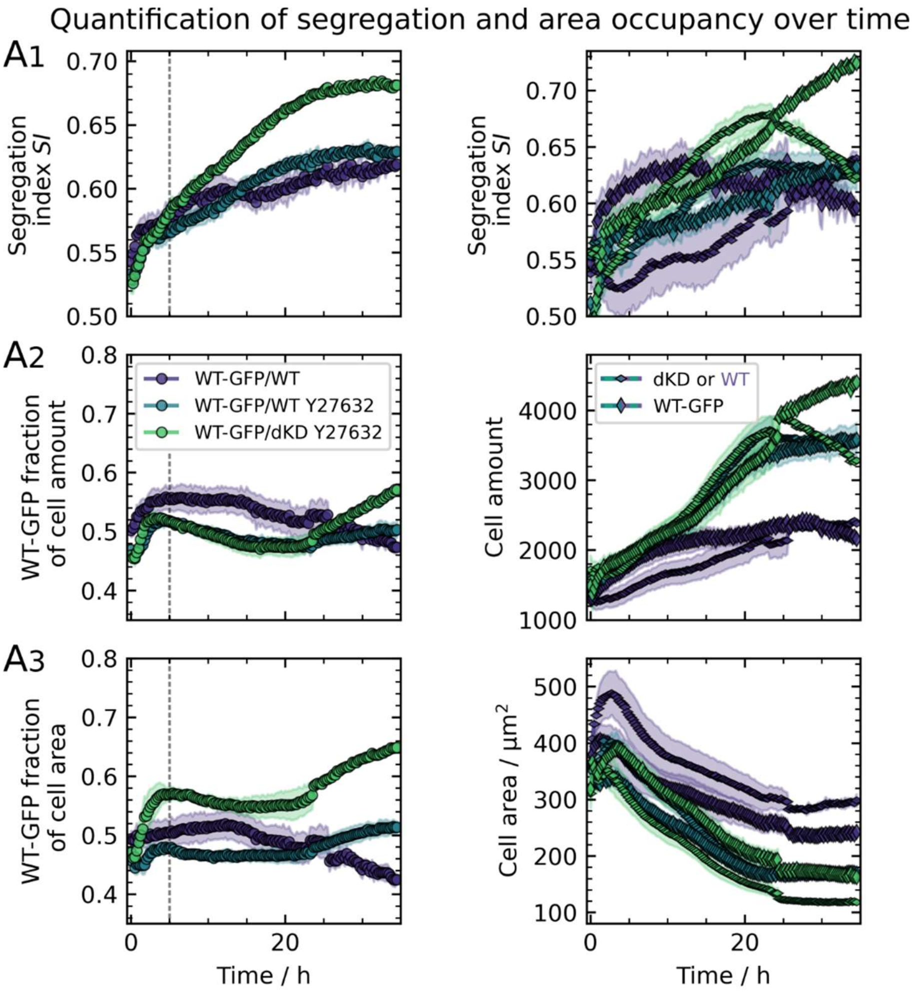


**Figure S8.** Contractility drives early, adhesion final sorting. Demixing behavior of dKD and wildtype cell co-cultures at an initial mixing ratio of 50:50, treated with 50 µM Y27632. All panels are set up as in Figure 5B and show the same data, and additionally untreated WT/WT from Figure 1 for comparison. Furthermore, longer experiment times are depicted, showing a plateau in the *SI* at the end of the experiment.

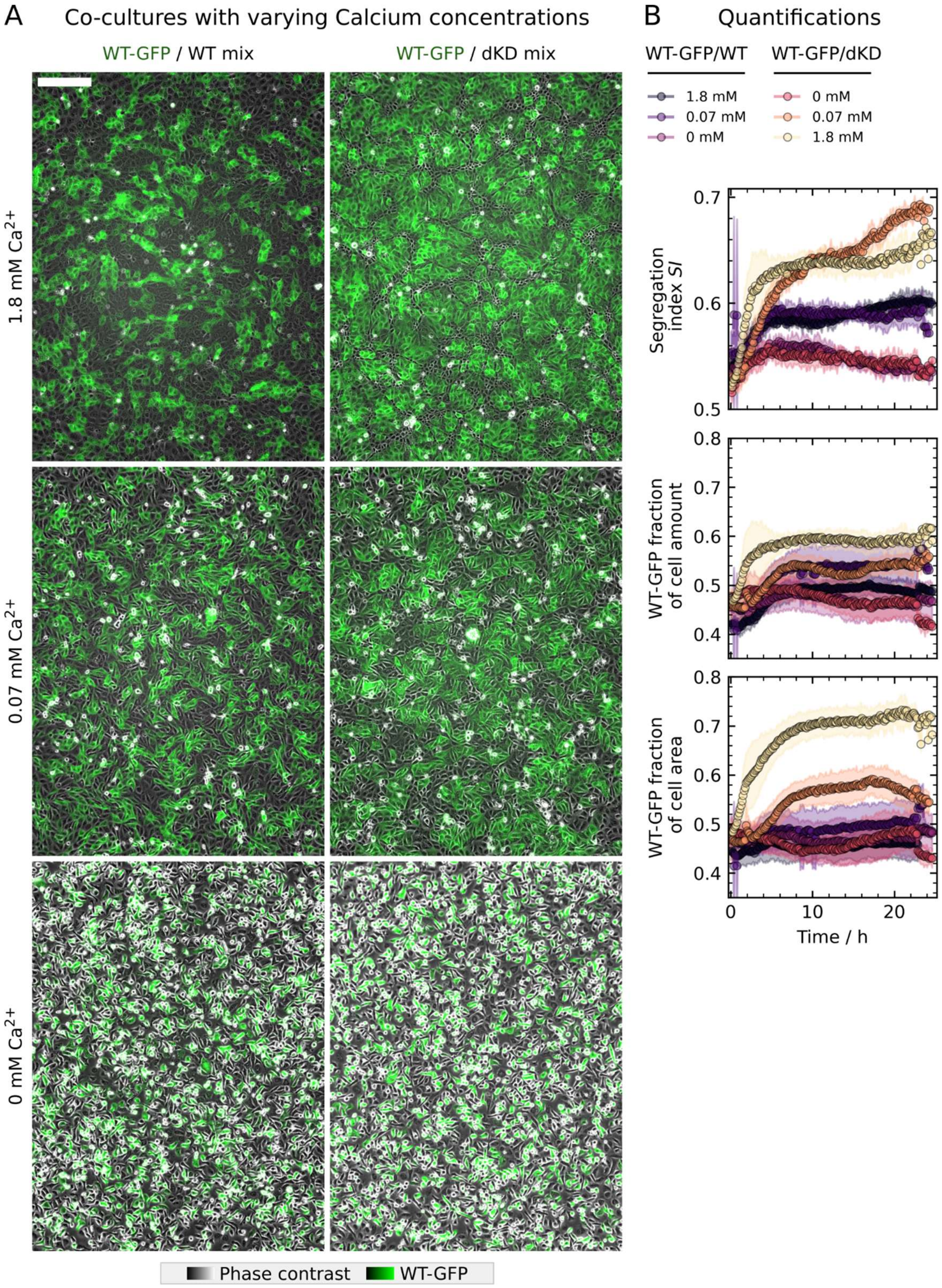


**Figure S9.** Calcium depletion abolishes both contractility and adhesion, and thereby also impairs demixing of dKD and WT cells. A) Example phase contrast images of co-cultures with superimposed WT-GFP fluorescence with varying calcium concentrations taken after 20 h of growth. B) Corresponding quantifications over time in accordance with the main Figures 1 and 5.

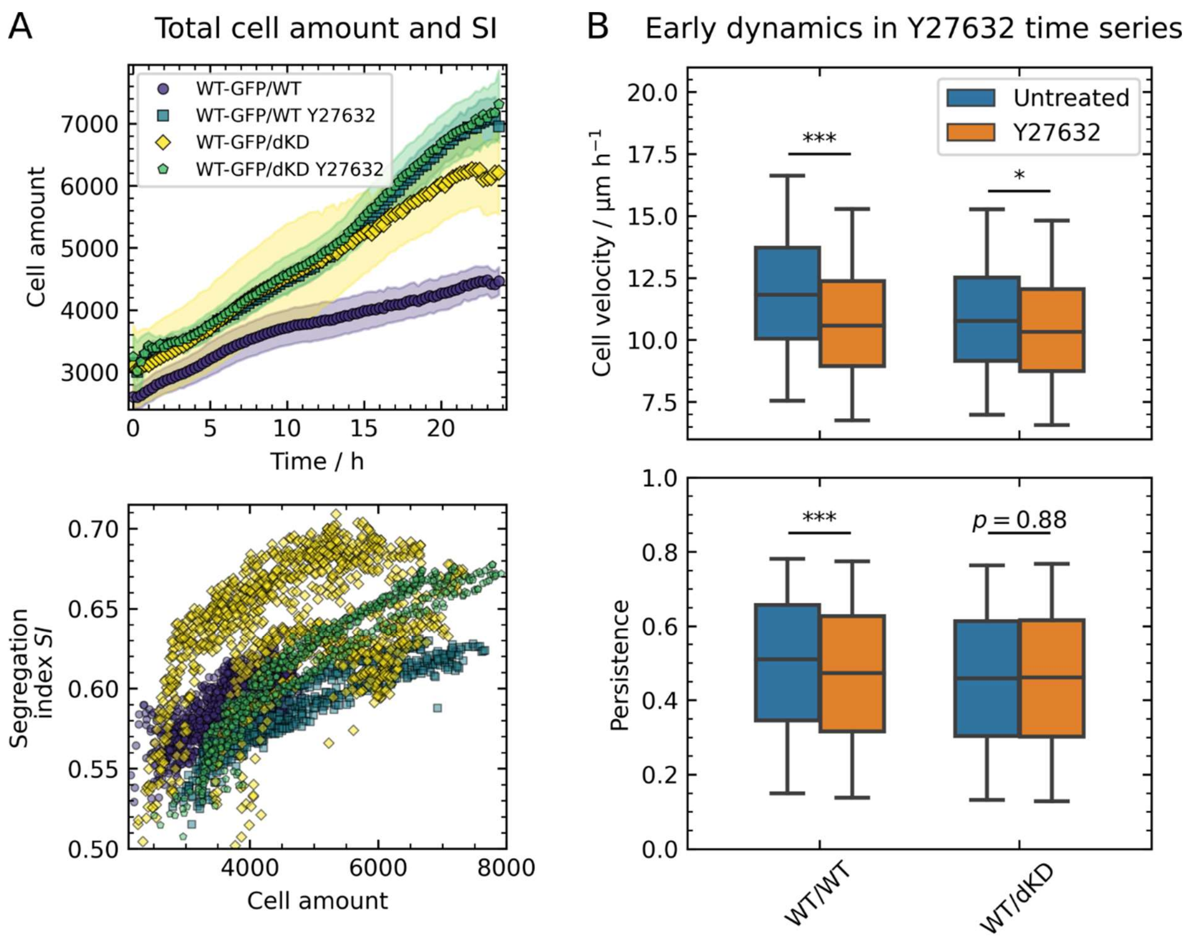


**Figure S10.** Additional analyses of the co-culture (de-) mixing experiments from Figures 1 and 5. A) Total cell amount over time and the *SI* plotted against the total cell amount. B) Velocity and persistence within the first 5 h after seeding. Both parameters were calculated from individual cell tracks. Velocity was averaged over all frames. Persistence was defined as the ratio of the direct distance between the start- and end points and the sum of the distances traveled in each step. Boxes show the median and upper and lower quartiles. Whiskers indicate the 5th and 95th percentile. The boxplot comprises individual cell velocity and persistence. For more accurate statistical testing, velocities and persistence were averaged over each separate time series.

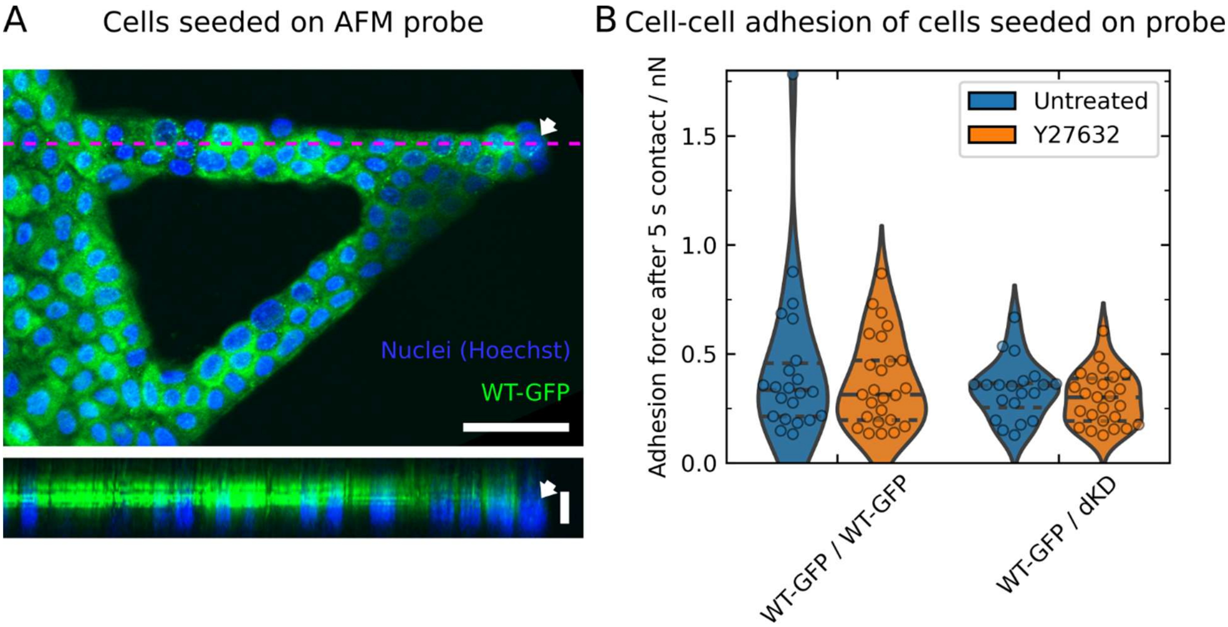


**Figure S11.** Cell-cell adhesion measurements with cells seeded on the backside of an AFM probe (cantilever). A) Confocal fluorescence image of cells adhered on an AFM probe and grown to confluence overnight. Arrows indicate the most outer cell, which will be the one in contact with a cell on the Petri dish during adhesion measurements. XY scale bar: 50 µm; Z scale bar: 15 µm. B) Quantification of adhesion forces after 5 s of contact between cells seeded on an AFM probe and subconfluent cells on a Petri dish. Violins represent a kernel density estimation with horizontal, dashed lines showing the quartiles and median. Violins are scaled to have the same area. Single data points represent individual adhesion peak forces. Three consecutive indentation/retraction cycles were performed for each cell pair. At least 8 cells on the substrate were measured, distributed over 2 days.
